# Multi-lab, Multi-enzyme Study Demonstrates the Versatility of Bacterial Microcompartment Shells as a Modular Platform for Confined Biocatalysis

**DOI:** 10.64898/2026.03.18.712704

**Authors:** Sreeahila Retnadhas, Nicholas M. Tefft, Yali Wang, Kyleigh L. Range, Arinita Pramanik, Kalpana Singh, Timothy K. Chiang, Kate Nigrelli, Robert P. Hausinger, Eric L. Hegg, Michaela A. TerAvest, Markus Sutter, Cheryl A. Kerfeld

## Abstract

Bacterial microcompartments (BMCs) are proteinaceous organelles that spatially organize metabolic reactions in bacteria and represent an attractive scaffold for pathway engineering. Here, we present a proof-of-concept *in vitro* study demonstrating a simple, scalable, and modular BMC shell-based platform for enzyme encapsulation using the SpyCatcher-SpyTag (SC-ST) covalent conjugation system. To evaluate the generality of this approach, 16 dehydrogenases were selected, of which 13 were successfully expressed and purified as SC-tagged enzymes in *E. coli* by five research groups working in parallel. Twelve of these efficiently conjugated to ST-fused BMC-T1 proteins, and addition of urea-solubilized BMC-H triggered rapid self-assembly of HT1 shells, resulting in successful encapsulation of all conjugated enzymes. The only enzyme lacking detectable activity after encapsulation was also inactive in its free SC-fused form, indicating that encapsulation retained enzymatic activity for all tested enzymes. Encapsulation modulated enzymatic activity and kinetic parameters in an enzyme-dependent manner, likely arising from variations in catalytic mechanism, structural flexibility affected by immobilization, and sensitivity to the local microenvironment created by encapsulation. Functional characterization of a subset of encapsulated enzymes revealed enhanced thermal stability up to ∼50 °C and improved storage stability relative to free SC-fused enzymes. Enzyme-loaded shells could be lyophilized and reconstituted without loss of structural integrity or activity. Finally, we demonstrate co-encapsulation of two enzymes within a single shell and their cooperative function through cofactor recycling. Together, these results establish engineered BMCs as a robust and modular platform for organizing multi-enzyme pathways, enabling rapid assembly, stabilization, and functional integration of enzymes for diverse metabolic engineering applications.

**Highlights:** A single strategy enables encapsulation of 12 diverse dehydrogenases in BMCs.

SpyCatcher-SpyTag interactions drive rapid enzyme assembly in BMCs.

Encapsulated enzymes are active and show improved thermal stability.

The platform enables scalable construction of synthetic metabolic modules.

**Graphical abstract:** 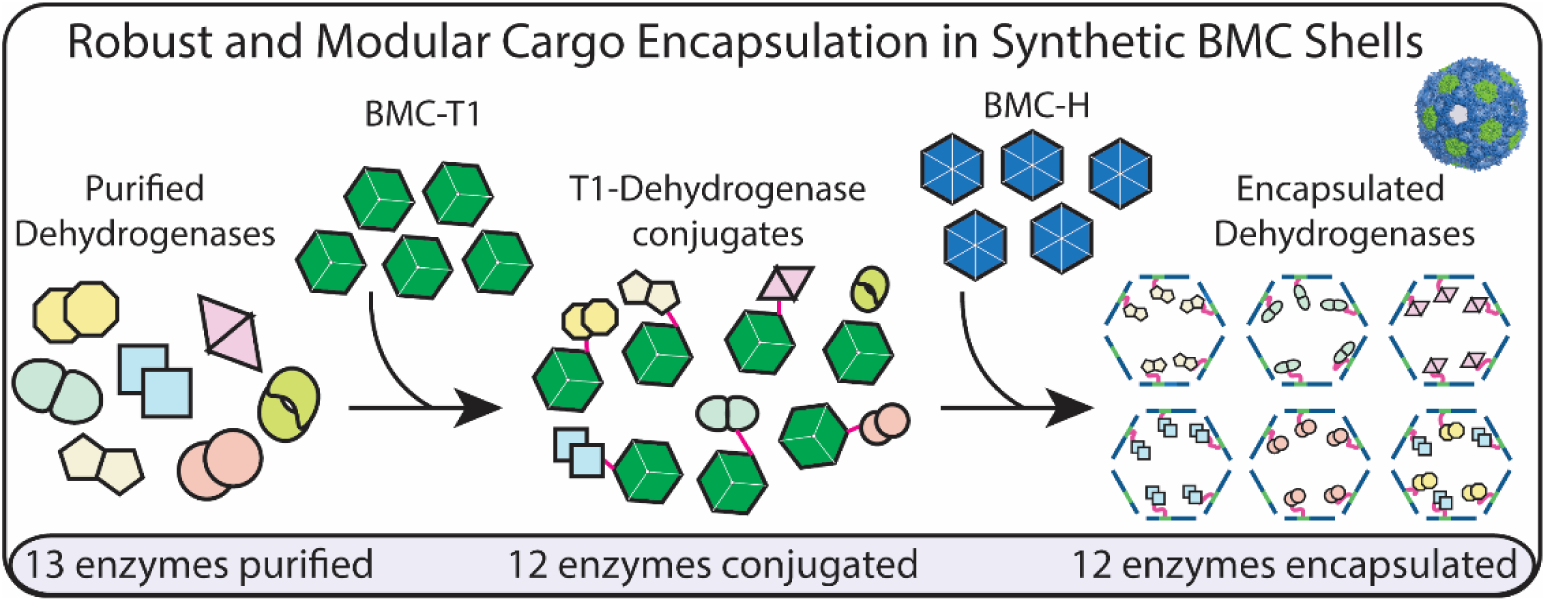

## 1. Introduction

Metabolic compartmentalization is a key innovation in the evolution of life (Ovádi and Saks, 2004). While the specialized functions of membrane-bound organelles of eukaryotes have long been understood, the appreciation of the extent of compartmentalization of catalytic reactions in bacteria is relatively recent. Based on comprehensive bioinformatic analyses, it is now known that many bacteria potentially form metabolic organelles known as bacterial microcompartments (BMCs) (Axen et al., 2014; Sutter et al., 2021; Sutter and Kerfeld, 2022). These are composed of a selectively permeable protein shell that surrounds segments of biochemical pathways. While there is recognizable homology among all BMC shell proteins, their functions are diverse, dictated by the encapsulated enzymes. There are at least 64 functionally distinct types of BMCs, including carboxysomes for CO_2_ fixation in cyanobacteria and some chemoautotrophs. However, most BMCs are catabolic, and these ‘metabolosomes’ share a core series of reactions (Axen et al., 2014; Kerfeld et al., 2018; Sutter et al., 2021) initiated by an aldehyde-generating enzyme referred to as the “signature” enzyme. The aldehyde is acted on by an NAD^+^ and CoA-dependent aldehyde dehydrogenase, forming NADH and an acyl∼CoA product. There are two known modes for regeneration of NAD^+^ in the lumen of BMCs, an encapsulated alcohol dehydrogenase that reduces a second aldehyde, forming an alcohol, or a recently identified microcompartment NADH dehydrogenase that recycles NAD^+^ via electron transport across the shell (Sutter et al., 2024).

While the signature enzymes and their cognate core enzymes vary widely, the shells of all BMCs are constructed from three classes of proteins. BMC-H proteins contain a single copy of the Pfam00936 domain and form hexamers, while BMC-T proteins are a fusion of two Pfam00936 domains that form trimers (pseudohexamers). BMC-H and BMC-T tiles constitute the facets of the polyhedral shell, while BMC-P proteins, composed of a single Pfam03319 domain, form pentamers that cap the vertices (Sutter et al., 2017; Greber et al., 2019; Sutter et al., 2019; Kalnins et al., 2020; Ferlez et al., 2023; Wang et al., 2024). Heterologous expression of shell proteins in the absence of cargo results in the formation of empty shells, providing the foundation for the design and engineering of synthetic BMCs. Numerous studies have demonstrated the potential to target non-native cargo such as the green fluorescent protein to these shells (Parsons et al., 2010; Choudhary et al., 2012; Lassila et al., 2014; Cai et al., 2016; Ferlez et al., 2023; Li et al., 2024). Functionally notable examples include bioreactors for ethanol (Lawrence et al., 2014) or hydrogen production (Li et al., 2020), formate utilization (Kirst et al., 2022), and the accumulation and storage of polyphosphate (Liang et al., 2017). These disparate studies collectively attest to the promise of BMC shells as a modular platform technology for catalysis in confinement.

To further advance the development of BMC shells as a generalizable technology for catalysis in confinement, we designed a multi-team experimental study to test the feasibility of loading a model BMC shell system with several members of a single class of enzymes. The model shell system is derived from the BMC found in *Haliangium ochraceum* (HO), which is extensively structurally characterized (Lassila et al., 2014; Sutter et al., 2017; Kirst et al., 2022) and for which orthogonal loading tags have been developed (Hagen et al., 2018a; Young et al., 2020). Moreover, there are *in vitro* assembly and rapid purification methods available (Hagen et al., 2018b; Range, Chiang et al., 2025) that, unlike heterologous expression and purification, provide greater control and simplify the assembly process. For our test case cargo, we selected a series of dehydrogenases (Fig. S1) including aldehyde dehydrogenase (EC 1.2.1.3), one of the most common enzyme classes associated with native BMC systems (Axen et al., 2014), which confine and process diverse aldehydes. Practically, the effects of tagging and encapsulation could be measured for all the enzymes by the same NAD(P)^+^ dependent assay, a component of the standardization of methods across different research groups.

Likewise, to control for variability in approach, the five research teams used a standardized shell system and loading protocol. We selected the HO HT1 “wiffle” shell, a 40 nm synthetic icosahedral shell that lacks pentamers at the vertices (Sutter et al., 2017; Hagen et al., 2018a; Kirst et al., 2022). While this shell form does not fully seal the compartment due to the ∼5 nm gaps at the vertices, it was chosen to remove the variable of differential substrate access to the encapsulated enzymes, which in a sealed shell relies substantially on access through the <10 Å pores in the BMC-H and BMC-T oligomers (Hagen et al., 2018a). HO HT1 shells can, if desired, readily be capped with HO BMC-P to form fully closed shells as previously demonstrated (Hagen et al., 2018a). As is, the HT1 shell can also be considered a three-dimensional scaffold on which to (co-)immobilize enzymes. Shells were assembled and loaded *in vitro* following a standardized formulation shared as a spreadsheet, and the effects of encapsulation on enzyme function were tested. For a subset of the designed shells, we further tested the effects of temperature and lyophilization on activity. We also constructed a two enzyme co-immobilized system that enabled local cofactor recycling. Using the same set of building blocks, members of the multi-laboratory team generated a diverse set of functionally active synthetic BMCs *in vitro* and in high yield using a standardized protocol for shell assembly and cargo loading (Fig. 1). Our results exemplify a generalizable, rapid prototyping method for constructing synthetic catalytic nanoreactors and provides a major advance in the development of BMC shells as a platform technology for catalysis in confinement.

**Fig. 1:**
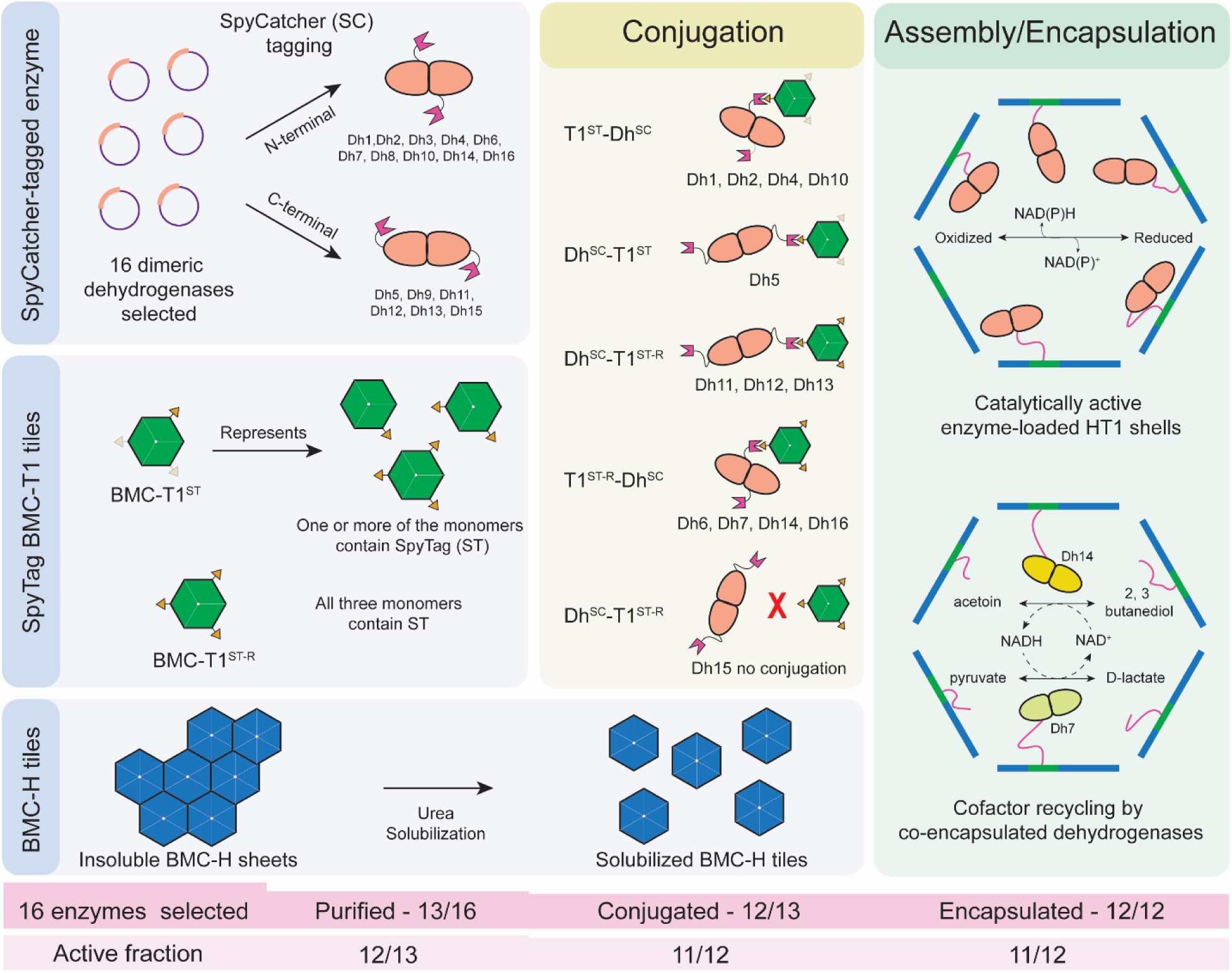
Workflow for encapsulating dehydrogenases within HT1 BMC shells: Encapsulation of dehydrogenases in HT1 shells was achieved using the SpyCatcher-SpyTag (SC-ST) system. **SpyCatcher-tagged enzyme:** Thirteen dehydrogenases were expressed in *E. coli* and purified by His_6_-affinity chromatography; eight were tagged with SC at the N-terminus and five at the C-terminus. Except for Dh12^SC^, all purified enzymes were catalytically active (12 of 13). **SpyTag BMC-T1 tiles:** Two versions of BMC-T1 tiles were used: BMC-T1^ST^ is a mixed tile containing at least one ST-bearing monomer per trimer, and BMC-T1^ST-R^ (replete) monomers all carry ST. **BMC-H tiles:** BMC-H proteins were purified as insoluble sheets and solubilized in 1.0-1.6 M urea to generate BMC-H tiles. **Conjugation block:** Except for Dh15^SC^, all dehydrogenases conjugated efficiently to both BMC-T1 tile variants via SC-ST interactions (12 of 13 enzymes), and all conjugates were catalytically active except Dh12^SC^ (11 of 12). **Encapsulation block:** Addition of BMC-H tiles to the conjugated trimer-enzyme complexes resulted in rapid assembly of HT1 shells, encapsulating all conjugated enzymes (12 of 12), with catalytic activity retained for all except Dh12^SC^ (11 of 12). Co-encapsulation of two dehydrogenases (Dh7^SC^ and Dh14^SC^) enabled mutual cofactor recycling, demonstrated by the formation of products from both enzymes upon addition of a single cofactor.

## 2. Methods

### 2.1 Cloning

The cloning strategies for dehydrogenases fused to either N- or C-terminal SpyCatcher003, for BMC-T1 tiles carrying both native BMC-T1 and SpyTagged-BMC-T1 (denoted as BMC-T1^ST^), and for BMC-T1 tiles carrying only SpyTagged-BMC-T1 (denoted as BMC-T1^ST-R^) are described in detail in the Supplementary Methods (Fig. S2). All DNA fragments were synthesized by Twist Bioscience with codon optimization for *E. coli*. Plasmid vectors were digested with restriction enzymes at 37 °C for 2 h, followed by alkaline phosphatase treatment for 1 h, preparative agarose gel electrophoresis, and gel extraction. Gene fragments were cloned into the vectors using Gibson assembly performed for 15 min at 50 °C, followed by transformation into *E. coli* DH5α cells and selection on LB agar plates containing kanamycin (50 µg/mL). Positive clones were identified by colony PCR and verified by Sanger sequencing. Sixteen of the seventeen dehydrogenase genes were successfully cloned into expression vectors.

### 2.2 Overexpression and purification

All dehydrogenases and both BMC-T1 variants (BMC-T1^ST^ and BMC-T1^ST-R^) were heterologously expressed in *E. coli* BL21(DE3) and purified by Ni²⁺-nitrilotriacetic acid (NTA) affinity chromatography followed by size-exclusion chromatography (SEC). Because experiments were performed by multiple researchers across different laboratories, expression conditions varied between enzymes and are described in detail in the supplementary methods. In general, protein expression was induced with anhydrotetracycline (aTc) for dehydrogenases and BMC-T1^ST^ constructs and IPTG for BMC-T1^ST-R^, followed by cell harvesting, lysis, Ni²⁺-affinity purification, and SEC.

BMC-H sheets were produced as described previously (Range and Chiang et al., 2025) with minor laboratory-specific modifications. Briefly, expression was induced with IPTG, cells were harvested and lysed, and BMC-H was recovered as insoluble sheets. The insoluble material was washed with detergent-containing buffer to remove cellular debris, followed by detergent-free washes, after which the BMC-H sheets were resuspended and aliquoted before freezing. Detailed lab-specific protocols are provided in the supplementary methods.

### 2.3 *In vitro* assembly

Aliquoted BMC-H sheets were prepared by reconstituting the insoluble sheet pellets in urea-containing TBS buffer (50 mM Tris-HCl, 150 mM NaCl, pH 8.0) and incubating at room temperature, followed by high-speed centrifugation to remove residual insoluble material. The urea concentration and incubation time varied between laboratories and are specified in the supplementary methods for the BMC-H used for encapsulating each dehydrogenase within HT1 shells. BMC-H concentrations were estimated by measuring absorbance at 280 nm and calculated using a theoretical extinction coefficient of 2,980 M⁻¹ cm⁻¹. *In vitro* assembly reactions (500 µL or 1 mL) were initiated by mixing BMC-T1^ST^ (or BMC-T1^ST-R^) and the respective dehydrogenase at defined monomer ratios in TBS supplemented with or without 10% (v/v) glycerol, followed by incubation at room temperature for a defined period to allow SpyTag-SpyCatcher conjugation. Detailed information is presented in the supplementary methods. BMC-H was then added at a BMC-H:BMC-T1 tile ratio of 3:1 to induce HT1 shell assembly. Reaction mixtures were clarified by high-speed centrifugation and applied to a size-exclusion chromatography column for shell purification. Assembled shells eluted in the void volume, whereas unincorporated components eluted according to their molecular sizes. Void-volume fractions were collected, and shell concentrations were determined using Pierce™ BCA Protein Assay Kits or absorbance at 280 nm measured by NanoDrop UV–vis (Thermo). Assembly quality was assessed by transmission electron microscopy (TEM) and dynamic light scattering (DLS).

Assembly of shells encapsulating two enzymes, HT1-(Dh7^SC^+Dh14^SC^), was performed as above except 550 µL of 1.84 mg/mL Dh14^SC^ and 225 µL of 3.6 mg/ml Dh7^SC^ was used to keep total cargo consistent with one enzyme shells.

### 2.4 Transmission Electron Microscopy

Glow-discharged 300-mesh Formvar-coated TEM grids (Ted Pella) were incubated with 10 μL of shell samples for 1 min, after which excess liquid was removed using Whatman filter paper. The grids were then rinsed two times with water, negatively stained with 10 μL of 1% uranyl acetate for 20-30 s, and blotted again to remove excess stain. Following air drying, the samples were imaged on a JEOL 1400 ‘Flash’ at 100 kV, and images were acquired using a Matataki Flash sCMOS camera.

### 2.5 Activity assays

The catalytic activity of each dehydrogenase was determined by continuously monitoring absorbance at 340 nm of the reaction mixture. Each enzyme reaction typically contained substrate, cofactor, and enzyme in buffer. Controls included reactions with (a) substrate and cofactor only (no protein), (b) with HT1-Dh^SC^ and substrate only (no cofactor), (c) with HT1-Dh^SC^ and cofactor only (no substrate), and (d) with HT1-empty shells, substrate, and cofactor. In oxidation reactions, reduction of NAD(P)⁺ to NAD(P)H caused an increase in A₃₄₀, whereas in reduction reactions, oxidation of NAD(P)H to NAD(P)⁺ caused a decrease in A₃₄₀. The slope of the linear portion of the absorbance curve was used to calculate enzyme activity using the extinction coefficient of NAD(P)H (6.22 × 10^3^ M^−1^ cm^−1^). Activity is reported as micromoles of NAD(P)H formed or consumed per min. Detailed protocols for each individual enzyme assay are provided in the supplementary methods. Dh16^SC^ activity was measured using the Glyceraldehyde 3 Phosphate Dehydrogenase Activity Assay Kit (Colorimetric) (Abcam, ab204732) as described in the product manual.

### 2.6 Assay for demonstrating cofactor recycling

Samples for testing two cargo BMCs were prepared in 2.0 mL glass HPLC vials (Vial: Restek, 21140; Cap: JG Finneran, 5395F09). 100 µL of 100 mM 2,3-butanediol, 100 µL of 100 mM sodium pyruvate, and 50 µL of 100 mM NADH or NAD^+^ were added. For conditions using purified enzymes, 2.5 µL of 3.6 mg/ml Dh7^SC^ and 5 µL of 1.8 mg/mL Dh14^SC^ were added. For conditions using HT1 shells 750 µL of 170 µg/mL purified shells were used. TBS 50/200 pH 8.0 was used to bring the final volume to 1 mL. Samples were incubated at 10 °C for 16 h prior to HPLC analysis.

HPLC analysis was performed on a Shimadzu 20A HPLC, using an Aminex HPX-87H (BioRad, Hercules, CA) column with a Microguard Cation H^+^ guard column (BioRad, Hercules, CA) at 60 °C. Compounds of interest were separated using a 0.6 mL/min flow rate, in 5 mM sulfuric acid with a 30 min run time. Eluent was prepared by diluting a 50% HPLC-grade sulfuric acid solution (Fluka) in water filtered through a 0.22 µm filter unit. Compounds of interest were detected by a refractive index detector (Shimadzu, RID-20A) maintained at 55 °C. Samples were prepared in 2.0 mL glass HPLC vials (Vial: Restek, 21140; Cap: Restek, 24485). Mixed standards of sodium pyruvate, sodium D-lactate, 2,3-butanediol, and acetoin were prepared at concentrations of 1, 2, 10, and 20 mM. Samples were maintained at 10 °C by an autosampler (Shimadzu, SIL-20AHT) throughout analysis. Sodium pyruvate, sodium D-lactate, 2,3-butanediol, and acetoin concentrations in the samples were determined using linear calibration curves based on the external standards.

## 3. Results

### 3.1 Dehydrogenases as Cargo Enzymes for BMC Shell Encapsulation via SpyTag-SpyCatcher

To demonstrate that BMC protein shells can be used for efficient encapsulation of a range of cargo enzymes while retaining catalytic activity upon encapsulation, a suite of structurally characterized dehydrogenases was selected (see Supplementary file 1). Dehydrogenases were chosen as the case-study enzyme class because BMCs in nature commonly sequester metabolic pathways that rely on dehydrogenases to process toxic and volatile aldehyde intermediates (Axen et al., 2014; Kerfeld et al., 2018). Additionally, these enzymes are all capable of being assayed spectroscopically by monitoring NAD(P)^+^ reduction or NAD(P)H oxidation despite the diversity of the catalyzed reactions, providing a single assay to be used across research teams. Specific selection criteria included an available structure in the Protein Data Bank (PDB), a functional annotation based on biochemical or *in vivo* characterization, and the commercial availability of substrates. We curated the list of candidates (Table 1) to ensure that we selected diverse activities that include reactions favoring both the NAD(P)H-oxidizing and NAD(P)^+^-reducing directions (Fig. S1); this duality facilitates downstream production of multi-enzyme cascade systems. We included variation in temperature optima: three of the selected enzymes (Dh11, Dh12, and Dh13) are derived from thermostable organisms, whereas the remaining enzymes originate from mesophiles. In this list of candidates, many enzymes have characteristics that may be useful in future applications.

**Table 1:** List of dehydrogenases used in this study.

| Name | Enzyme (annotated as) | SpyCatcher (SC) tagging | Monomeric MW with SC tag<sup>a</sup> | EC number | PDB | Oligomeric state<sup>b</sup> | Source organism | Substrate tested | Comments |
| --- | --- | --- | --- | --- | --- | --- | --- | --- | --- |
| **Dh1** | Aldehyde reductase | N-terminal | 56 kDa | 1.1.1.1 | 4QGS | Dimer | *Escherichia coli* | Propionaldehyde | Encapsulated and active in HT1 |
| **Dh2** | Alcohol dehydrogenase | N-terminal | 54 kDa | 1.1.1.1 | 3OX4 | Dimer | *Zymomonas mobilis* | Ethanol | Encapsulated and active in HT1; Lyophilization compatibility studied |
| **Dh3** | Glycerate dehydrogenase | N-terminal | 49 kDa | 1.1.1.29 | 1GDH | Dimer | *Hyphomicrobium methylovorum* | Not tested | Poor expression |
| **Dh4** | Acetaldehyde dehydrogenase | N-terminal | 68 kDa | 1.2.1.5 | 1EYY | Dimer | *Vibrio harveyi* | Butyraldehyde | Encapsulated and active in HT1; Storage stability studied |
| **Dh5** | Propionaldehyde dehydrogenase | C-terminal | 61 kDa | 1.2.1.10 | 4C3S | Dimer | *Clostridium phytofermentans* | Propionaldehyde | Encapsulated and active in HT1; Storage stability studied |
| **Dh6** | L-Lactate dehydrogenase | N-terminal | 51 kDa | 1.1.1.27 | 4UUL | Dimer | *Trichomonas vaginalis* | Pyruvate | Encapsulated and active in HT1<sup>R</sup>; Substrate kinetics and thermal stability studied |
| **Dh7** | D-Lactate dehydrogenase | N-terminal | 50 kDa | 1.1.1.28 | 7JP2 | Dimer | *Treponema pallidum* | Pyruvate | Encapsulated and active in HT1<sup>R</sup>; Substrate kinetics and thermal stability studied; Demonstrated cofactor recycling with Dh14<sup>SC</sup> |
| **Dh8** | Malate dehydrogenase | N-terminal | 48 kDa | 1.1.1.37 | 4TVO | Dimer | *Mycobacterium tuberculosis* | Not tested | Poor expression |
| **Dh9** | Aminomuconate-semialdehyde dehydrogenase | C-terminal | Not cloned | 1.2.1.32 | 4I25 | *Dimer | *Pseudomonas fluorescens* | Not tested | Failed cloning |
| **Dh10** | Leucine dehydrogenase | N-terminal | 54 kDa | 1.4.1.9 | 8GXD | *Dimer | *Exiguobacterium sibiricum* | Leucine | Encapsulated and active in HT1 |
| **Dh11** | L-Lactate dehydrogenase | C-terminal | 48 kDa | 1.1.1.27 | 1Y6J | *Dimer | *Clostridium thermocellum* | L-lactate | Encapsulated and active in HT1<sup>R</sup> |
| **Dh12** | Malate dehydrogenase | C-terminal | 53 kDa | 1.1.1.37 | 1V9N | Dimer | *Pyrococcus horikoshii* | L-lactate/L-malic acid/oxaloacetate | Encapsulated in HT1<sup>R</sup>, but no detectable activity even for free Dh12<sup>SC</sup> |
| **Dh13** | Malate dehydrogenase | C-terminal | 49 kDa | 1.1.1.37 | 1IZ9 | Dimer | *Thermus thermophilus* | L-Malic acid | Encapsulated and active in HT1<sup>R</sup> |
| **Dh14** | Butanediol dehydrogenase | N-terminal | 51 kDa | 1.1.1.4 | 6IE0 | Dimer | *Bacillus subtilis* | 2, 3-butanediol | Encapsulated and active in HT1<sup>R</sup>; Thermal stability studied; Demonstrated cofactor recycling with Dh7<sup>SC</sup> |
| **Dh15** | Formate dehydrogenase | C-terminal | 54 kDa | 1.17.1.9 | 8HTY | Dimer | *Candida boidinii* | Formate | Did not conjugate to T1-ST<sup>R</sup> |
| **Dh16** | Glyceraldehyde 3-phosphate dehydrogenase | N-terminal | 49 kDa | 1.2.1.12 | 1DC3 | Dimer | *Escherichia coli* | Glyceraldehyde 3-phosphate | Encapsulated and active in HT1<sup>R</sup> |

<sup>a</sup>Theoretical monomeric molecular weight of Dh<sup>SC</sup> (Dh fused with SC) calculated using ExPASy ProtParam from the sequences in their respective plasmid constructs
<sup>b</sup>Oligomeric states as reported in their respective PDB entries
\*Surface area buried when crystallographic symmetry is expanded suggests functional dimeric states while they are annotated as monomers (Dh10 and Dh11) or tetramer (Dh9) in the PDB

To target the dehydrogenase enzymes to the shell lumen, we used a previously developed strategy where a SpyTag was inserted in a lumen facing loop of BMC-T1 (Hagen et al., 2018a) that would autocatalytically crosslink with a SpyCatcher modified cargo protein. To increase conjugation efficiency, we used the recently developed SpyCatcher003-SpyTag003 (SC-ST hereafter) system (Keeble et al., 2019). A His_6_-tagged SC was attached to the cargo enzymes at either the N- or C-terminus with a short (GS)_3_ linker after consideration of the available PDB structures to select termini that are solvent-exposed and unlikely to impede oligomerization or activity; if purification tags were used in the crystallized protein, we chose, for example, the His_6_-tagged terminus as the site of the SC fusion.

### 3.2 Robust Production of BMC Shell Components for HT1 Assembly

To highlight the robustness of this strategy, the protein purifications, conjugations, encapsulations, and activity assays were performed in parallel by seven researchers across five laboratories at two institutions. Of the 16 enzymes selected for the study, we were unable to clone the gene encoding one enzyme (Dh9), and two SC fused enzymes did not express well (Dh3 and Dh8). The other thirteen SC-tagged dehydrogenases (denoted Dh^SC^) were heterologously expressed and purified following the protocols described in the Materials and Methods.

Structural components of the HT1 shell, BMC-H and BMC-T1^ST^ (ST fused to BMC-T1), were also expressed and purified as described in the Materials and Methods by each laboratory. Two different BMC-T1^ST^ constructs were used in this study. To prevent formation of aggregates from crosslinking of the oligomeric cargo and shell proteins (Fig. S2A), we designed a construct carrying both native BMC-T1 and BMC-T1^ST^ (Fig. S2B) (such tiles denoted as BMC-T1^ST^). These tiles contain at least one ST monomer for enzyme conjugation (because only the tagged copy is His_6_-tagged), but judging from sodium dodecylsulfate-polyacrylamide gel electrophoresis (SDS-PAGE), about 1-2 untagged copies were present. This construct was used for the assembly of Dh1^SC^, Dh2 ^SC^, Dh4^SC^, Dh5^SC^, and Dh10^SC^. For the assembly of the other dehydrogenases (Dh6^SC^, Dh7^SC^, Dh11^SC^, Dh12^SC^, Dh13^SC^, Dh14^SC^, Dh15^SC^, and Dh16^SC^), a construct carrying only BMC-T1^ST^ was employed (Fig. S2B). Because all three monomers in the trimer have SpyTag fusions, we designate these latter tiles as “replete” denoted BMC-T1^ST-R^ (Fig. S2A, Fig. S2B). The choice of type of BMC-T1 tile used was arbitrary, selected by the individual investigator; our objective was to be able to compare assembly competence and activity between loaded shells based on both types of trimers.

Optimized methods (Range, Chiang et al., 2025) for the expression and purification of BMC-H sheets resulted in a yield of ∼150 mg per liter of culture, which could be used for ∼300 individual 500-µL HT1 shell assembly reactions (corresponding to a total of ∼200 mg of assembled HT1 shells). Similarly, single-step Ni²⁺-affinity purification of BMC-T1^ST^ and BMC-T1^ST-R^ produced a similar yield of ∼10 mg per liter of expression culture, which could be used for ∼40 individual 500-µL shell assembly reactions (corresponding to a total of ∼26 mg of assembled HT1 shells). All shell components can be flash frozen in aliquots without affecting assembly efficiency. All laboratories involved in the study were able to efficiently express and purify the HT1 shell components with high yield and purity as shown in Fig. S3.

### 3.3 Dehydrogenases are efficiently loaded into HT1 shells using SC-ST

Cargo loading of dehydrogenases into HT1 protein shells was achieved via conjugation of Dh^SC^ to BMC-T1^ST^ (or BMC-T1^ST-R^) trimer tiles. SDS-PAGE analysis indicated that SC-ST conjugation was highly efficient for all tested enzymes (except Dh15^SC^), as evidenced by high molecular weight bands corresponding to the combined size of BMC-T1^ST^ (or BMC-T1^ST-R^) and Dh^SC^ observed in SDS-PAGE gels (Fig. S3). Interestingly, two prominent bands were detected for some conjugated dehydrogenases in SDS-PAGE gels (Fig. S3G). This doublet was most apparent for shells containing BMC-T1^ST-R^, where all three monomeric units carry the ST. Mass spectrometric analysis of excised bands, however, confirmed the presence of BMC-T1 and the corresponding dehydrogenases in both of these higher molecular weight species, indicating that they could represent species that have different migration due to distinctions arising from the crosslink.

An IVA (*in vitro* assembly) worksheet was designed and used to standardize and simplify calculation of the HT1 shell components and cargo proteins required for shells with varying cargo loading and easy scale-up (Supplementary file 2). Empty HT1 shells were assembled using a molar tile ratio of 3:1 for BMC-H to BMC-T1, a ratio that reflects the ratio in the final shell structure and that was previously shown to be optimal for shell formation (Range, Chiang et al., 2025). Consistent with these reports, efficient assembly was observed using this ratio, as confirmed by size exclusion chromatography (SEC) (Fig. 2M) and transmission electron microscopy (TEM) analyses (Fig. S4A). For dehydrogenase-loaded HT1 shells, both types of BMC-T1 tiles (BMC-T1^ST^ and BMC-T1^ST-R^) assembled efficiently into HT1 shells, yielding enzyme-loaded shells reproducibly across different laboratories (Fig. 2A-K, Fig. S3 and Fig. S4). SEC confirmed the assembly of individual components into protein shells (Fig. 2), with TEM images showing well-formed, uniform shells (Fig. S4). Three monomeric ratios of BMC-T1^ST^ (BMC-T1^ST-R^) to dehydrogenase were used for encapsulation of these enzymes into HT1 shells - 3:1 (Fig. 2A), 3:2 (Fig. 2F, Fig. 2H-L), and 3:3 (Fig. 2B-D, Fig. 2G). Under these conditions, near-complete encapsulation was observed for all dehydrogenases except Dh6^SC^, as indicated by negligible SEC peaks corresponding to SC-tagged free or conjugated enzymes (Fig. 2A-K). This uniformity demonstrates that both the crosslinking reaction as well as shell assembly are highly efficient and complete despite adding relatively bulky cargo to the shell proteins.

**Fig. 2.**
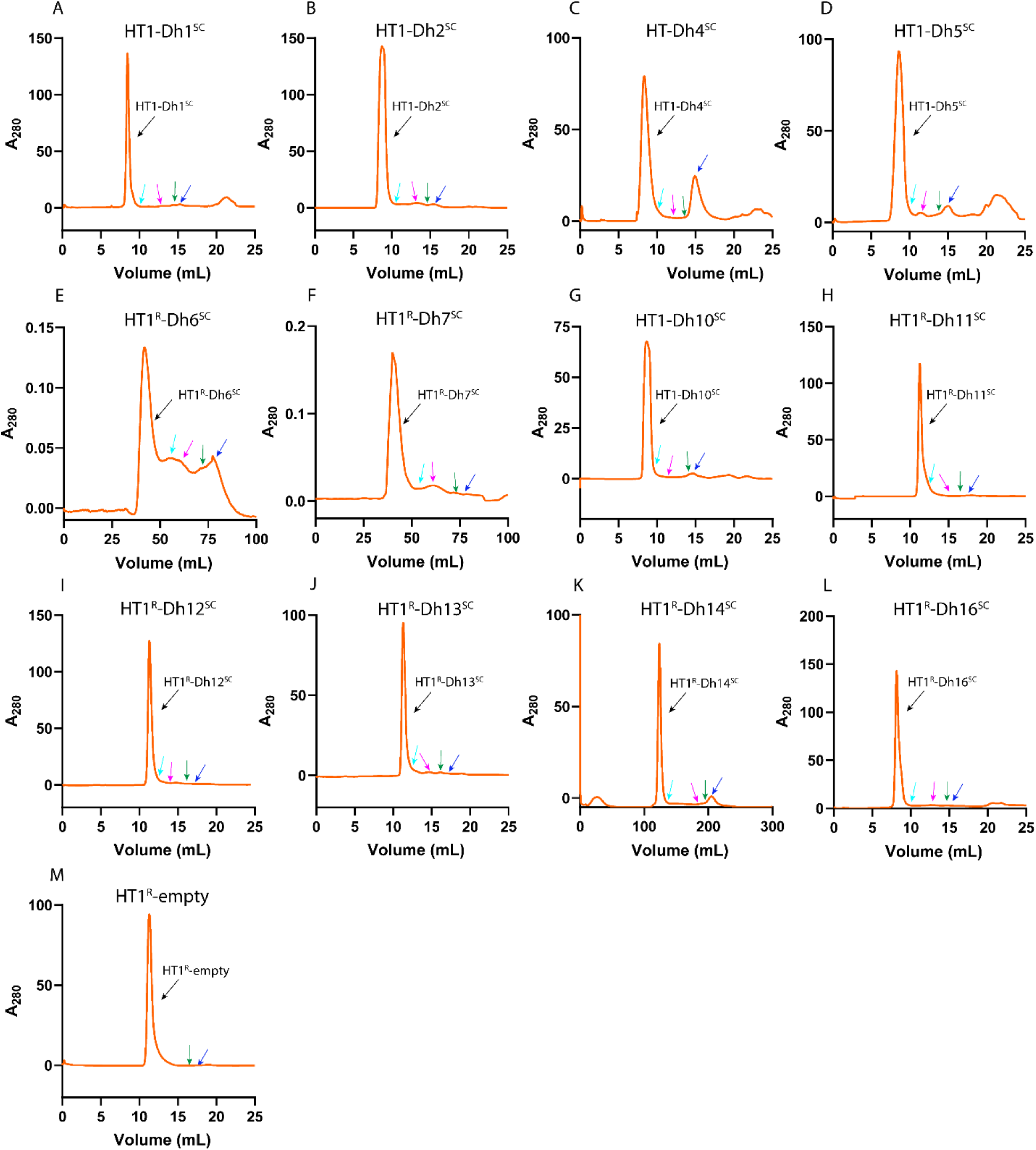
Size-exclusion chromatography (SEC) purification of *in vitro* assembly (IVA) reactions for empty and cargo-loaded HT1 shells. For IVA reactions, purified dehydrogenases (Dh^SC^) and BMC-T1^ST^ (or BMC-T1^ST-R^) were mixed at a monomeric ratio of either 1:3 (A), 2:3 (E, F, G), or 3:3 (B, C, D) and incubated at room temperature for 30-60 min prior to the addition of BMC-H. BMC-H was added at a tile ratio of 3:1 (BMC-H:BMC-T1^ST^), which immediately initiated HT1 shell assembly. Due to the rapidity of assembly, reaction mixtures were filtered or centrifuged at high speed to remove aggregates and immediately injected onto a Superdex 200 column. Assembled HT1 shells eluted in the void volume (as marked in the figure), while unincorporated components were separated based on size. The presence of a single dominant peak in the void volume, with no significant peaks corresponding to free Dh^SC^ (marked as 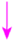), BMC-T1^ST^ or BMC-T1^ST-R^ (marked as 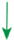), BMC-H (marked as 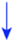), or Dh-trimer conjugates (marked as 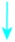), indicates efficient assembly and near-complete incorporation of cargo enzymes. HT1^R^ (E, F, H, I, J, K, L) denotes HT1 shells containing BMC-T1^ST-R^ tiles whereas HT1 (A, B, C, D, G) denotes HT1 shells containing BMC-T1^ST^ tiles.

### 3.4 Differential behavior of dehydrogenases upon conjugation and encapsulation

#### 3.4.1 All tested dehydrogenases retain activity upon conjugation and subsequent encapsulation

Catalytic activities of the dehydrogenases (except Dh16^SC^) were measured spectroscopically by monitoring absorbance at 340 nm (A₃₄₀), corresponding to the formation or consumption of NAD(P)H during substrate oxidation or reduction reactions, respectively. Dh16^SC^ activity was measured using a commercially available glyceraldehyde-3-phosphate dehydrogenase activity assay kit, which monitors the reaction by measuring absorbance at 450 nm (A₄₅₀). Activities were measured for the free SC-tagged enzyme (Dh^SC^), the trimer-conjugated enzyme (Dh^SC^-T1^ST^), and the HT1-encapsulated enzyme (HT1-Dh^SC^) as described in the methods section. With one exception, the enzymes retained at least some catalytic activity upon conjugation to both types of BMC-T1 trimer tiles and after subsequent assembly into HT1 shells (Fig. 3). For Dh12^SC^, which is annotated as a malate dehydrogenase, no catalytic activity was detected with L-malate, oxaloacetate, pyruvate, or L-lactate as substrates. Despite the absence of any detectable activity under the conditions tested, Dh12^SC^ conjugated efficiently to BMC-T1^ST-R^ tiles and assembled into HT1 shells (Fig. 2I), underscoring the robustness and modularity of the HT1 shell assembly system. This, and the observation that 12 of 13 enzymes retained activity when encapsulated highlights the ease and general applicability of the HT1 platform for enzyme loading, enabling reliable shell assembly across diverse enzymes; these initial prototypes are poised for optimization.

**Fig. 3.**
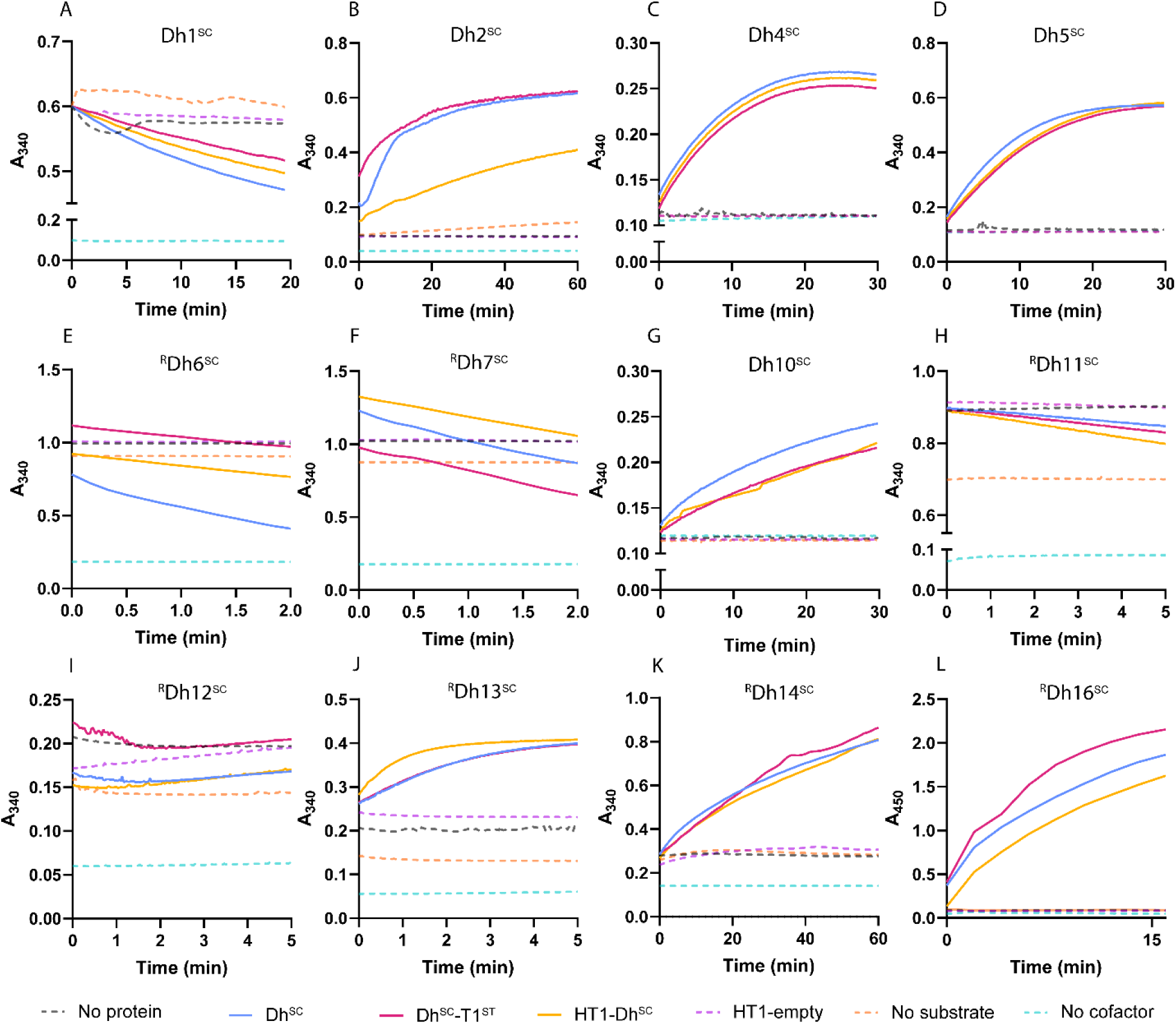
Catalytic activities of dehydrogenases upon conjugation to BMC-T1^ST^ and subsequent encapsulation into HT1 shells. Each panel shows the catalytic activity of a different Dh^SC^ (labeled on figure), measured by continuously monitoring absorbance at 340 nm (A₃₄₀). For Dh16^SC^, a commercially available kit was used to measure its activity by monitoring absorbance at 450 nm (A_450_). In reactions where the substrate is oxidized, NAD(P)⁺ is reduced to NAD(P)H, resulting in an increase in A₃₄₀ (B, C D, G, I, J and K) as the reaction progressed. Conversely, in reactions where the substrate is reduced, NAD(P)H is oxidized to NAD(P)⁺, resulting in a decrease in A₃₄₀ (A, E, F and H). However, for Dh12^SC^, the minimal increase in A_340_ in the presence of any protein component, including the HT1-empty control, suggests that there is no significant activity from the enzyme component. Control reactions are shown as dashed lines, and reactions containing free Dh^SC^, BMC-T1^ST^ conjugated Dh^SC^, or encapsulated Dh^SC^ are shown as solid lines. BMC-T1^ST^ tiles were used for the assembly of Dh1^SC^, Dh2^SC^, Dh4^SC^, Dh5^SC^, and Dh10^SC^, while BMC-T1^ST-R^ tiles were used for the assembly of Dh6^SC^, Dh7^SC^, Dh11^SC^, Dh12^SC^, Dh13^SC^, and Dh14^SC^. The data shown are representative traces from one of three independent experiments performed for each enzyme.

Because all tested dehydrogenases except Dh6^SC^ showed nearly complete enzyme incorporation into HT1 shells (Fig. 2A-K), the catalytic activity was normalized per milligram of enzyme (Table 2). This calculation was based on the known molar ratios of dehydrogenase to BMC-T1^ST^ (or BMC-T1^ST-R^) used in the assembly mixture, assuming complete incorporation of the dehydrogenase into the HT1 shells (Fig. S5).

**Table 2:** Effect of conjugation and encapsulation on enzyme activity.

| Dehydrogenase | BMC-T1<sup>ST</sup>:Dh<sup>SC</sup><br>(Monomers) | Dh<sup>SC</sup><br>monomers<br>per HT1 | Specific activity |  |  |
| --- | --- | --- | --- | --- | --- |
|  |  |  | Dh<sup>SC</sup> % | Dh<sup>SC</sup>-BMC-T1<br>% | HT1-Dh<sup>SC</sup><br>% |
| Dh1<sup>SC</sup> | 3:1 | 20 | 100 ± 4 | 65 ± 2 | 77 ± 2.7 |
| Dh2<sup>SC</sup> | 3:3 | 60 | 100 ± 24 | 81 ± 11 | 17 ± 0.3 |
| Dh4<sup>SC</sup> | 3:1 | 60 | 100 ± 3 | 87 ± 4 | 93 ± 2.5 |
| Dh5<sup>SC</sup> | 3:1 | 60 | 100 ± 4 | 98 ± 3 | 96 ± 4 |
| <sup>R</sup>Dh7<sup>SC</sup> | 3:2 | 40 | 100 ± 5 | 100 ± 11 | 90 ± 10 |
| Dh10<sup>SC</sup> | 3:3 | 40 | 100 ± 5 | 97 ± 21 | 87 ± 20 |
| <sup>R</sup>Dh11<sup>SC</sup> | 3:2 | 40 | 100 ± 8 | 124 ± 7 | 128 ± 4 |
| <sup>R</sup>Dh13<sup>SC</sup> | 3:2 | 40 | 100 ± 10 | 100 ± 13 | 124 ± 19 |
| <sup>R</sup>Dh14<sup>SC</sup> | 3:2 | 40 | 100 ± 14 | 88 ± 10 | 62 ± 7 |
| <sup>R</sup>Dh16<sup>SC</sup> | 3:2 | 40 | 100 ± 5 | 86 ± 10 | 131 ± 8 |

<sup>R</sup>Conjugation and subsequent encapsulation were performed using BMC-T1<sup>ST-R</sup>

Dh1^SC^ and Dh4^SC^ showed a partial reduction in activity upon conjugation, but exhibited increased activity after encapsulation, whereas Dh2^SC^ and Dh14^SC^ exhibited small losses in activity upon conjugation with substantial decreases following encapsulation (Table 2, Fig. S5A-B, Fig. S5C, Fig. S5I). Interestingly, two enzymes Dh11^SC^ and Dh16^SC^ displayed a small yet significant increase in activity following conjugation and encapsulation (Fig. S5G and Fig. S5J), while other enzymes like Dh5^SC^, Dh7^SC^, Dh10^SC^ and Dh13^SC^ showed little, if any, change in activity under either condition (Table 2, Fig. S5D-F, Fig. S5H).

Under the tested conditions, no consistent trend in enzyme activity was observed following conjugation or encapsulation between enzymes conjugated to the different BMC-T1 tiles (BMC-T1^ST^ and BMC-T1^ST-R^). Likewise, no trend was observed associated with the different Dh^SC^ loading ratios tested (Table 2).

Similarly, encapsulation of enzymes within HT1 shells modulated their kinetic behavior in an enzyme-dependent manner. For Dh7^SC^, encapsulation resulted in a pronounced decrease in *K*_m_, from approximately 123 μM for the free SC-tagged enzyme to 47 μM for HT1-Dh7^SC^, corresponding to a ∼2.6-fold increase in apparent substrate affinity, with only a moderate reduction in *V*_max_ from 11.8 to 8.1 μmol min⁻¹ (Fig. S6A-C). In contrast, Dh6^SC^ exhibited a substantial reduction in *V*_max_ accompanied by an increase in *K*_m_ upon encapsulation, indicating a stronger kinetic penalty (Fig. S6D-F). Collectively, our results demonstrate that the impact on catalytic activity of conjugation and subsequent encapsulation into HT1 seems to be enzyme dependent, but almost all enzymes retain activity, underscoring the potential of the HT1 shell as a generic platform for confining enzymatic cargo.

#### 3.4.2 HT1 shell encapsulation differentially impacts dehydrogenase stability

Additional tests of the effects of encapsulation were performed for a subset of the dehydrogenases. Thermal stability assays further revealed that shell encapsulation conferred clear stabilization effects, particularly for Dh6^SC^, with encapsulated HT1-Dh6^SC^ maintaining significantly greater residual activity at elevated temperatures (40 °C to 50 °C) compared to the free SC-tagged enzyme (Fig. S7A). This enhanced thermal stability, however, was accompanied by a marked reduction in catalytic throughput for Dh6^SC^ as discussed in the previous section, reflecting a common trade-off between stabilization and maximal reaction rate upon encapsulation, potentially due to restricted flexibility and altered substrate access. Dh7^SC^ appears to be intrinsically more thermally stable than Dh6^SC^, but at elevated temperatures, encapsulated HT1-Dh7^SC^ retained slightly greater relative activity than Dh7^SC^. Nevertheless, both Dh7^SC^ and HT1-Dh7^SC^ completely lost activity above 50 °C under the conditions tested, suggesting limits to shell-mediated protection at higher temperatures (Fig. S7B). For Dh14^SC^, the stabilization conferred by encapsulation was intermediate, being less pronounced than for Dh6^SC^ and greater than for Dh7^SC^, with a steady decrease in activity as temperature increased and encapsulation provided a significant enhancement in stability at all temperatures below 52 °C (Fig. S7C). Collectively, these results indicate that shell encapsulation can improve enzymatic robustness under mild heat stress but does not universally prevent thermal inactivation at more extreme temperatures.

Stability studies upon storage of Dh4^SC^ and Dh5^SC^ conducted over a five-week period showed that the free SC-tagged enzyme, T1-conjugated enzyme, and shell-encapsulated enzyme did not exhibit any loss of activity when stored at 4 °C. In contrast, encapsulation into HT1 shells markedly improved storage stability at room temperature. After two weeks of storage at room temperature, Dh4^SC^ and BMC-T1-Dh4^SC^ retained ∼30% of their day-0 activity, whereas HT1-Dh4^SC^ retained ∼80% (Fig. S8A). Similarly, Dh5^SC^ and BMC-T1-Dh5^SC^ retained ∼10-15% of their day-0 activity after 2 weeks, while HT1-Dh5^SC^ retained ∼65% (Fig. S6B) of its day-0 activity. By the end of five weeks at room temperature, all forms of Dh4^SC^ and Dh5^SC^ had completely lost activity. Nevertheless, encapsulation (HT1-Dh^SC^) consistently conferred greater stability compared to the free SC-tagged enzymes (Dh^SC^) until week 4. These results demonstrate that encapsulation substantially improves storage stability under conditions in which the free enzyme is unstable.

We also tested if catalytically active shells could be lyophilized with catalytic activity retained upon re-hydration. Assembled HT1-Dh2^SC^ shells tolerated lyophilization without any significant loss of catalytic activity (Fig. S9A). TEM images of encapsulated HT1-Dh2^SC^ before and after lyophilization confirmed that the structural integrity of the shells was well preserved (Fig. S9B), consistent with the retention of Dh2^SC^ activity.

While all but one enzyme retained activity upon encapsulation, the specific effects of SpyTag-mediated conjugation and encapsulation within HT1 shells on enzymatic activity are enzyme-specific. Although encapsulation within HT1 confers thermal stabilization, the extent of this effect varies depending on the intrinsic stability and properties of each enzyme. Furthermore, encapsulation also appears to improve stability upon storage at room temperature and loaded shells are amenable to lyophilization for long term storage.

### 3.5 Co-encapsulation of two enzymes demonstrates functional cofactor recycling between enzymes

To evaluate the ability for co-encapsulation of multiple enzymes and to demonstrate cofactor recycling, Dh7^SC^ (D-lactate dehydrogenase) and Dh14^SC^ (2,3-butanediol dehydrogenase) were co-encapsulated within HT1 shells by performing *in vitro* assembly of the respective T1-linked enzymes (Fig. S10A) at an approximately equal ratio which resulted in shells with negligible leftover unassembled proteins (Fig. S10B). While we cannot guarantee that the resulting individual shells have equal amounts of each enzyme, the stochastic distribution makes a similar amount likely. TEM confirmed intact and uniform shell formation (Fig. S10C). Both enzymes retained catalytic activity after encapsulation, indicating that HT1 assembly preserves individual enzyme function. Dh7^SC^ exhibits substantially greater catalytic activity than Dh14^SC^ under the tested conditions, rendering Dh14^SC^ rate limiting and thereby enabling clear assessment of cofactor coupling.

Cofactor recycling between Dh7^SC^ and Dh14^SC^ was evaluated using two complementary reaction conditions designed to test enzyme coupling under defined redox constraints (Fig. 4A and 4B). In the first condition (Fig. 4A), 5 mM NADH was supplied as the sole cofactor in reactions containing 10 mM pyruvate, 10 mM 2,3-butanediol, and co-encapsulated enzyme shells, HT1-(Dh7^SC^+Dh14^SC^). Under these conditions, Dh7^SC^ rapidly reduced pyruvate to D-lactate, oxidizing NADH to NAD⁺ (Fig. 4C). After overnight incubation at 10 °C, the co-encapsulated enzyme shells produced approximately 5.22 mM D-lactate, slightly exceeding the initial NADH concentration. The accumulation of ∼0.8 mM acetoin (Fig. 4D) demonstrates that Dh14^SC^ utilized the NAD⁺ generated by Dh7^SC^ to oxidize 2,3-butanediol, thereby regenerating ∼0.8 mM NADH. This regenerated NADH supported the additional pyruvate reduction by the faster Dh7^SC^. However, because NADH regeneration was constrained by the slower catalytic rate of Dh14^SC^, NADH availability ultimately limited further pyruvate reduction, resulting in incomplete conversion of pyruvate to D-lactate (Fig. 4A). These results confirm Dh14^SC^ as the rate-limiting step in the cofactor regeneration cycle.

**Fig. 4:**
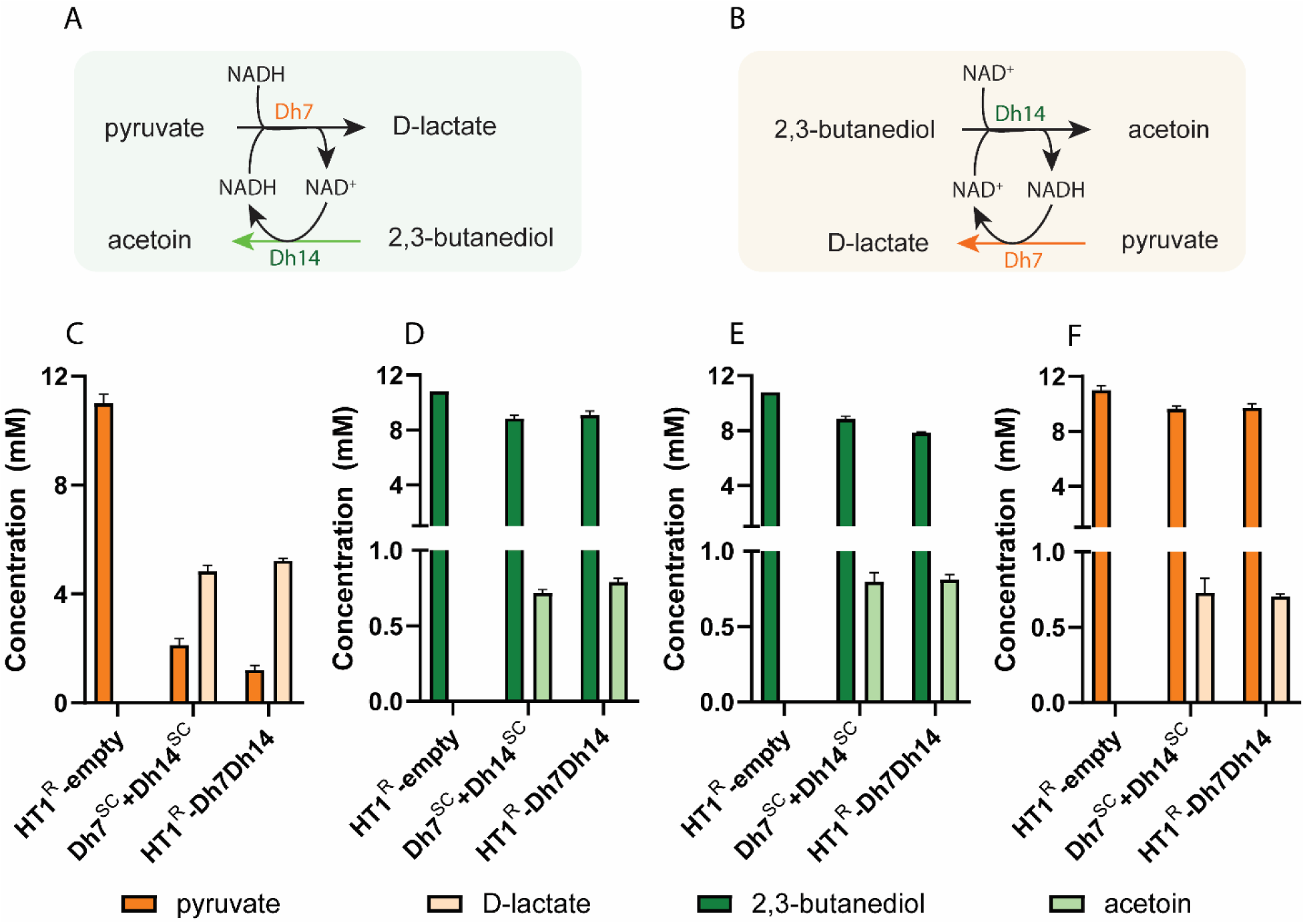
Cofactor recycling by co-encapsulated Dh7^SC^ and Dh14^SC^. Cofactor recycling by Dh7^SC^ and Dh14^SC^ was evaluated under two complementary reaction conditions. **A)** A reaction in which NADH was supplied as the sole cofactor; NADH was oxidized by Dh7^SC^ during the reduction of pyruvate to lactate, generating NAD⁺, which in turn supported the catalytic oxidation of 2,3-butanediol to acetoin by Dh14^SC^. **B)** A complementary reaction in which NAD⁺ was supplied as the sole cofactor under otherwise identical conditions. In this case, catalytic reduction of pyruvate to D-lactate by Dh7^SC^ depends on the generation of NADH by Dh14^SC^ during the oxidation of 2,3-butanediol. Concentrations of reactants and products for **C)** Dh7^SC^ catalyzed reaction (pyruvate and D-lactate) and **D)** Dh14^SC^ catalyzed reaction (2,3-butanediol and acetoin) when the reaction contained both substrates (pyruvate and 2,3-butanediol) and both enzymes (Dh7^SC^ and Dh14^SC^), either as free SC-tagged enzymes or co-encapsulated within HT1 shells, with NADH as the sole cofactor. Concentrations of reactants and products for **E)** Dh14^SC^ catalyzed reaction (pyruvate and D-lactate) and **F)** Dh7^SC^ catalyzed reaction (2,3-butanediol and acetoin) under identical reaction conditions but with NAD⁺ as the sole cofactor.

This conclusion was further supported by experiments in which identical reactions were performed with NAD⁺ as the sole cofactor (Fig. 4B). In this configuration, Dh7^SC^ activity was strictly dependent on NADH generated by Dh14^SC^. Dh14^SC^ oxidized 2,3-butanediol to produce ∼0.8 mM acetoin (Fig. 4E), and the resulting NADH enabled Dh7^SC^ to generate ∼0.7 mM D-lactate from pyruvate (Fig. 4F). The close correspondence between acetoin formation and D-lactate production demonstrates direct coupling between the two reactions and confirms that Dh14^SC^ controls the overall flux through the cofactor regeneration cycle. For comparison, a positive control reaction containing free Dh7^SC^ and Dh14^SC^ exhibited similar product profiles to the encapsulated system, whereas a negative control containing HT1-empty shells showed no detectable product formation. This outcome shows that co-encapsulated Dh7^SC^ and Dh14^SC^ function cooperatively within HT1 shells to regenerate redox cofactors in a spatially confined environment.

These results demonstrate that HT1 shells can co-encapsulate multiple enzymes while preserving individual catalytic activities and enabling cooperative function. This approach is analogous to the functional organization observed in natural bacterial microcompartments (Chowdhury et al., 2014). Together, these findings support a mechanistic model in which engineered protein shells like HT1 not only provide a structural scaffold for enzyme encapsulation but also create a localized microenvironment that enhances coupled catalysis and can support multienzyme cascades, mirroring natural microcompartment function.

## 4. Discussion

Cofactor recycling and metabolite channeling by the co-encapsulated core enzymes of BMCs is one of the many attributes that have made them the focus of numerous efforts to develop them as nanobioreactors (Gonzalez-Esquer et al., 2016; Plegaria and Kerfeld, 2018; Doron et al., 2023; Li et al., 2024; Shinde and Chowdhury, 2024; Johnson et al., 2025; Snyder et al., 2025). Other possible attributes of BMCs include isolation of toxic or inhibitory/volatile intermediates, sequestration of enzymes from competing reactions, increased stability of enzymes due to immobilization, and the selective permeability of the shell that also provides the potential for establishing specialized microenvironments for catalysis. The strategy we used here for a standardized design, build, and test approach for bottom-up assembly of synthetic BMCs, carried out in parallel by five research groups at 2 different sites, was remarkably successful. Robust shells were generated *in vitro* rapidly and efficiently (with only trace amounts of unassembled building blocks) and loaded with cargo; among the 13 different enzymes expressed and purified from *E. coli*, 12 conjugated efficiently to SpyTag003-fused BMC-T1 (with the only exception being Dh15^SC^), and all conjugates assembled efficiently into wiffle-ball-like HT1 shells.

Apart from one enzyme (Dh12), for which activity could not be detected even in the free SpyCatcher-fused form (Dh12^SC^), all enzymes retained catalytic activity after conjugation and encapsulation. Upon encapsulation, one enzyme (Dh2^SC^) exhibited a dramatic reduction in activity (∼80%), four enzymes (Dh5^SC^, Dh7^SC^, Dh10^SC^, and Dh13^SC^) showed no significant change, two enzymes (Dh11^SC^ and Dh16^SC^) displayed a mild increase in activity (∼10%), two enzymes (Dh1^SC^ and Dh14^SC^) showed a moderate reduction in activity (30–40%), and one enzyme (Dh4^SC^) showed only a small reduction (∼5%). For Dh6^SC^, accurate estimation of the encapsulated enzyme fraction was not possible because not all the enzyme present in the reaction was encapsulated within the shells; nevertheless, measurable catalytic activity was retained following encapsulation.

While all enzymes tested retained activity upon encapsulation, the extent varied. The decrease in apparent *K*_m_ for HT1-Dh7^SC^ may reflect changes in the microenvironment or restricted conformational flexibility upon encapsulation, which can stabilize catalytically competent conformations and lower the energetic barrier for substrate binding (Ahmed et al., 2019; Ibrahim et al., 2016; Liu et al., 2024). In HT1-Dh6^SC^, the increased *K*_m_ (Figure S4D-F) likely reflects reduced substrate accessibility at the active site, potentially due to unfavorable enzyme orientation or microenvironmental effects imposed by the shell (Arana-Peña et al., 2021). The relatively modest reduction in *V*_max_ suggests that shell confinement does not strongly limit turnover for Dh7^SC^, indicating favorable compatibility with the HT1 shell.

The observed variability in activity among different enzymes likely reflects differences in their intrinsic catalytic mechanisms and structural flexibility—influenced by immobilization and sensitivity to local microenvironmental changes imposed by encapsulation. Several non-mutually exclusive factors may contribute to these differences. First, the relative positioning of the active site with respect to the shell interior and the SpyCatcher fusion site may influence substrate accessibility and catalytic turnover. Second, enzyme-shell interactions may further modulate activity. Enzyme immobilization has been shown to impact activity and the HT1 shells are effectively a three-dimensional scaffold upon which the dehydrogenases were tethered. Furthermore, electrostatic, hydrophobic, and/or transient physical interactions between the cargo enzyme and HT1 shell proteins could perturb local folding and alter active-site geometry. Such effects are expected to vary depending on enzyme surface properties and charge distribution, providing a plausible explanation for the enzyme-dependent trends observed here. Third, encapsulation also imposes a confined and crowded microenvironment, which can differentially affect enzymatic function. Molecular crowding has been shown to stabilize folded states and suppress aggregation, thereby enhancing enzyme stability and lifetime (Cheung et al., 2005; Gimón et al., 2025). However, crowding can simultaneously restrict conformational fluctuations required for catalysis (Ma and Nussinov, 2013) and can contribute to reduced activity upon encapsulation. Finally, it is likely that the availability of substrate to shell-immobilized enzymes is less than it is for an enzyme free in bulk solution, an inherent tradeoff for a system that relies on selective permeability to substrates and products for an overall enhancement of catalysis.

Moreover, kinetic and stability studies performed on a subset of these enzymes indicate that encapsulation within HT1 can confer thermal stabilization; the extent of this effect varies depending on the intrinsic stability and other properties of each enzyme.

## 5. Conclusion

In summary, we have developed a design, build, test workflow for precisely assembling synthetic BMCs for rapid prototyping of nanoreactors for catalysis in confinement. Key features of our approach include a standardized formula for constructing shells from their building blocks using a defined, orthogonal method for loading cargo (Spy-Tag/SpyCatcher) and *in vitro* assembly which obviates the time-intensive process of tuning (co-)expression conditions and the subsequent purification of assembled shells for assays. By systematically evaluating a broad panel of dehydrogenases, we demonstrated that this approach supports efficient cargo loading, robust shell assembly, and retention of catalytic activity across multiple enzymes. Encapsulation conferred measurable improvements in both thermal and storage stability of cargo enzymes. We further demonstrate cooperative activity between two encapsulated enzymes through efficient cofactor recycling within the confined environment. Collectively, these findings establish the HT1 BMC shell system as a scalable and versatile platform for organizing catalysts and constructing synthetic nanoreactors. It not only provides a structural scaffold for catalyst encapsulation and immobilization, but it can also create a localized microenvironment that enhances coupled catalysis through controlled cofactor recycling, mimicking natural BMC function.

## CRediT authorship contribution statement

**Sreeahila Retnadhas:** Writing – original draft, Writing – review and editing, Data curation, Formal analysis, Investigation, Methodology, **Nicholas M. Tefft:** Formal analysis, Investigation, Methodology, **Yali Wang:** Formal analysis, Investigation, Methodology, **Kalpana Singh:** Formal analysis, Investigation, Methodology, **Arinita Pramanik:** Formal analysis, Investigation, Methodology, **Kyleigh L. Range:** Formal analysis, Investigation, Methodology, **Tim K. Chiang:** Investigation, **Kate Nigrelli:** Investigation, **Robert Hausinger:** Funding acquisition, Supervision, Validation, Writing – review and editing, **Eric L. Hegg:** Funding acquisition, Supervision, Validation, Writing – review and editing, **Michaela A. TerAvest:** Funding acquisition, Supervision, Validation, Writing – review and editing, **Markus Sutter:** Validation, Writing – review and editing, Formal analysis, Investigation, Methodology, and **Cheryl A. Kerfeld:** Funding acquisition, Supervision, Validation, Writing – original draft, Writing – review and editing

## Conflict of interest

The authors declare no conflict of interest

## Acknowledgements

Research was supported as part of the Center for Catalysis in Biomimetic Confinement, an Energy Frontier Research Center funded by the U.S. Department of Energy (DOE), Office of Science, Basic Energy Sciences (BES), under award DE-SC0023395.

## Supplementary files

Supplementary file 1: Supplementary text, methods and figures (.docx file)

Supplementary file 2: *In vitro* assembly (IVA) worksheet (.xlsx file)

## Data availability

Data will be made available on request.

## Notes

### Competing Interest Statement

The authors have declared no competing interest.

