## Supplemental file 1 for "Multi-lab, Multi-enzyme Study Demonstrates the Versatility of Bacterial Microcompartment Shells as a Modular Platform for Confined Biocatalysis"

1. **Summary of all enzyme examples**

Dh1 is the very well characterized zinc- and $NADP^+$-dependent alcohol dehydrogenase from *Escherichia coli* (YqhD, EC 1.1.1.1, PDB: 1OJ7) that exhibits a broad substrate range (Fig. S1A) and has proven useful for production of renewable biofuels and chemicals (Sulzenbacher et al., 2004; Jarboe, 2011). Dh2 is the iron- and $NAD^+$-dependent alcohol dehydrogenase from *Zymomonas mobilis* (EC 1.1.1.1, PDB: 3OX4) that exhibits greatest specificity for ethanol/acetaldehyde (Fig. S1A, where R = methyl) (Moon et al., 2011). Dh3 is the $NAD^+$-dependent D-glycerate dehydrogenase (Fig. S1B) from the obligate methylotrophic bacterium *Hyphomicrobium methylovorum* (EC 1.1.1.29, PDB: 1GDH) (Goldberg et al., 1994). Dh4 is the $NADP^+$-dependent acetaldehyde dehydrogenase (Fig. S1C) required for bioluminescence in the bacterium *Vibrio harveyi* (EC 1.2.1.5, PDB: 1EYY) (Ahvazi et al., 2000). Dh5 is the $NAD^+$-dependent propionaldehyde dehydrogenase from the cellulolytic soil bacterium *Lachnoclostridium* (formerly *Clostridium*) *phytofermentans* (EC 1.1.1.27, PDB: 4C3S) that is naturally found in a bacterial microcompartment and uses coenzyme A to form the thioester product (Fig. S1D) (Tuck et al., 2016). Dh6 is an $NAD^+$-dependent L-lactate dehydrogenase (Fig. S1E) from *Trichomonas vaginalis* (EC 1.1.1.27, PDB 4UUL), the flagellated protozoan parasite of humans (Steindel et al., 2016). Dh7 is the $NAD^+$-dependent D-lactate dehydrogenase (Fig. S1F) from the microaerophilic, spirochete *Treponema pallidum* responsible for syphilis in humans (EC 1.1.1.28, PDB: 7JP2) (Deka et al., 2020). Dh8 is the $NAD^+$-dependent L-malate dehydrogenase (Fig. S1G) from the human pathogen *Mycobacterium tuberculosis* that is responsible for tuberculosis (EC 1.1.1.37, PDB: 4TVO) (Ferraris et al., 2015). Dh9 is the $NAD^+$-dependent 2-aminomuconate-semialdehyde dehydrogenase (Fig. S1H) used in the degradation pathways of 2-aminophenol in *Micromonospora rosaria* (Doron et al., 2023) and 2-nitrobenzoic acid in *Pseudomonas fluorescens* (EC 1.2.2.32, PDB 4I25) (Huo et al., 2015); however, the substrate of this protein is not available due to its cyclization to form picolinic acid thus diminishing interest in the enzyme. Dh10 is an $NAD^+$-dependent leucine dehydrogenase (Fig. S1I) from *Exiguobacterium sibiricum* (EC 1.4.1.9, PDB: 8GXD) (Zhao et al., 2025). Enzymes from thermophilic microorganisms are of special interest for biotechnology, so Dh11, Dh12, and Dh13 are additional $NAD^+$-dependent dehydrogenases of L-lactate, malate, and malate from the thermophiles *Clostridium thermocellum* (PDB: 1Y6J), *Pyrococcus horikoshii* (PDB: 1V9N), and *Thermus thermophilus* (PDB: 1IZ9) (unpublished). Dh14 is the $NAD^+$-dependent 2R,3R-butanediol dehydrogenase (Fig. S1J) from *Bacillus subtilis* (EC 1.1.1.4, PDB: 6IE0) (unpublished). Dh15 is an $NAD^+$-dependent formate dehydrogenase (Fig. S1K) from *Candida boidinii* (EC 1.17.1.9, PDB: 8HTY) (Gul et al., 2023). Finally, Dh16 is the $NAD^+$-dependent glyceraldehyde 3-phosphate dehydrogenase (Fig. S1L) from *E. coli* (EC 1.2.1.12, PDB: 1DC3) (Yun et al., 2000).

2. **Construct considerations and cloning strategies**

In the natural BMC system, the cargo is thought to be targeted to the inside of the shells with non-covalent encapsulation peptides that bind to the interior shell surface. In our synthetic system, we use a covalent SpyTag003/SpyCatcher003 (Keeble et al., 2019) encapsulation method with the SpyTag on an internal loop of the BMC-T1 shell protein (Hagen et al., 2018) and SpyCatcher on the cargo for localization to the shell interior. However, the covalent nature of the crosslink can hinder proper shell assembly when multimeric proteins (as dehydrogenases usually are) linked to different T1 shell proteins (Fig. S2A), thereby creating a chain of crosslinked BMC-T1-cargo protein assemblies and could cause aggregation and prevent efficient shell formation.

To address this issue, we designed a construct that can reduce the number of BMC-T1$^{ST}$ protomers and thereby reduce the amount of cargo localization. In addition to the BMC-T1 incorporating a SpyTag003 on an inside facing loop flanked by GGGGS-GGGGS linkers we added a second wild type BMC-T1 copy(no His-tag and no SpyTag) (Fig. S2B). After co-expression and purification, we then obtained a mixture of BMC-T1 with at least one but up to three SpyTags. Band intensity on sodium dodecylsulfate-polyacrylamide gel electrophoresis (SDS-PAGE) gels (Supplementary Fig. S3A) suggests there is a significant amount of wild type BMC-T1, we estimate that most BMC-T1 proteins contain only one or two SpyTags. We refer to this version as BMC-T1$^{ST}$ and the version that has a SpyTag on all copies as BMC-T1$^{ST\text{-}R}$.

We used two different strategies for N- vs C-terminal tagging with SpyCatcher003. For the N-terminally tagged versions (10 dehydrogenases) we constructed a pBbE2K-based vector encoding a Met-6xHis-GS-SpyCatcher003-GSGSGS protein sequence that is followed by a BamHI restriction site. The dehydrogenase genes were designed with 5' and 3' adapters to allow Gibson assembly into the BamHI-cut vector. Additionally, the 3' adapter included a sequence element for an additional ribosome binding site and BamHI site. This procedure allows adding extra copies of dehydrogenase genes without a SpyCatcher tag (Fig. S2C).

For the C-terminally tagged versions (7 dehydrogenases) this strategy is not feasible due to the repeat elements causing issues with gene synthesis, so we made constructs that already included SpyCatcher003 and that were cloned using Gibson assembly into an NdeI/BamHI-cut pBbE2K vector (Fig. S2D).

Initial tests with conjugation indicated good assembly even with dehydrogenases that have two SpyCatchers, so we did not need to clone versions that express mixed tagged/untagged dehydrogenases. However, this step could be useful for enzymes that form higher order oligomers than dimers. Both BMC-T1$^{ST}$ and BMC-T1$^{ST\text{-}R}$ BM were successfully used for conjugating dehydrogenases as well as forming shells so the reduction of the number of SpyTags did not seem to have a significant effect. This could be due to steric clashes where the attachment of one cargo molecule already prevents addition of more cargo even if there are more SpyTags available.

### 3. Supplementary methods:

#### 3.1 Overexpression and purification

**Dh1:** Dh1 with an N-terminal $His_6$-SpyCatcher003 fusion ($Dh1^{SC}$) and mixed BMC-T1 tiles (BMC-$T1^{ST}$) were heterologously expressed in *E. coli* BL21(DE3) and purified by $Ni^{2+}$-nitrilotriacetic acid (NTA) affinity chromatography followed by size-exclusion chromatography. Plasmids were transformed into *E. coli* BL21(DE3), and a 50 mL culture inoculated with multiple colonies was grown overnight at 37 °C in LB containing 50 µg/mL kanamycin . One liter of LB supplemented with kanamycin was inoculated with 2% (v/v) overnight culture and grown at 37 °C, 180 rpm until the $OD_{600}$ reached 0.6-0.8. Expression was induced with 100 ng/µL anhydrotetracycline (aTc), and cultures were grown overnight at 22 °C. Cells were harvested, flash-frozen, and stored at -80 °C. Cell pellets were resuspended in 40 mL TBS (50 mM Tris-HCl, pH 8.0, 150 mM NaCl) containing 20 mM imidazole and lysed using a French Press at 25,000 psi. After centrifugation at 20,000 × *g* for 45 min, the soluble fraction was applied to a 5 mL $Ni^{2+}$-NTA HiTrap column, washed with 20 mM imidazole and 300 mM NaCl, and eluted with 300 mM imidazole in TBS. Eluted proteins were concentrated and further purified by size-exclusion chromatography (SEC) on a Superdex™ 200 column equilibrated in TBS. Purified proteins were concentrated, aliquoted and either flash-frozen and stored at -80 °C (BMC-$T1^{ST}$) or on ice ($Dh1^{SC}$).

BMC-H sheets were overexpressed by inoculating 1 L LB medium containing 50 µg/mL ampicilin with 2% (v/v) overnight culture and growing at 37 °C, 180 rpm to an $OD_{600}$ of 0.6-0.8. Expression was induced with 0.2 mM IPTG, and cultures were incubated overnight at 22 °C with shaking at 180 rpm. Cells were harvested, flash-frozen, and stored at -80 °C. BMC-H sheet purification was performed as described in our previous study (Range, Chiang et al., 2025). Frozen pellets were resuspended in 30 mL hexamer lysis buffer (50 mM Tris-HCl pH 8.0, 100 mM NaCl, 10 mM $MgCl_2$) supplemented with DNase I and lysozyme and lysed using a French Press. Triton X-100 was added to 1% (v/v), and lysates were incubated for 20 min at room temperature with gentle agitation. Insoluble white material enriched in BMC-H sheets formed on top of cell debris was collected after centrifugation at 20,000 rpm for 20 min, washed repeatedly with hexamer wash buffer (containing 1% Triton X-100), followed by a final wash without detergent. The final pellet was resuspended in 10 mL hexamer lysis buffer, aliquoted, pelleted and stored at -80 °C as aliquots.

**Dh2 and Dh10:** *E. coli* BL21(DE3) chemically competent cells (New England Biolabs) were transformed with plasmid DNA containing $Dh2^{SC}$, $Dh10^{SC}$, mixed BMC-T1 (BMC-$T1^{ST}$), or BMC-H protein sequences according to the vendor's specifications. Cells were incubated with 50–100 ng of plasmid DNA on ice for 30 min and heat shocked at 42 °C for 10 s. Cells were then placed back on ice for 5 min before adding 950 µL of Luria–Bertani (LB) broth for 1 h at 37 °C with gentle shaking. 100 µL of recovered cells was plated on kanamycin ($Dh2^{SC}$ , $Dh10^{SC}$, and BMC-$T1^{ST}$) or ampicillin (BMC-H) plates, and colonies were grown overnight at 37 °C. Colonies of transformed BL21(DE3) cells were grown in 50 mL of LB broth with 50 µg/mL kanamycin ($Dh2^{SC}$ $Dh10^{SC}$, and BMC-$T1^{ST}$) or 100 µg/mL ampicillin (BMC-H) at 37 °C overnight. Cells were induced at OD 0.6–0.8 at 600 nm with 50 µL of 1 mg/mL aTc ($Dh2^{SC}$, $Dh10^{SC}$, and BMC-$T1^{ST}$) or 500 µL of 1 M IPTG (BMC-H) per 1 L of culture. The induced cells were incubated at

22 °C for 18 h. The cells were pelleted by centrifugation at 8000 rpm for 20 min at 4 °C with a JLA 8.1 rotor in a Beckman Coulter Avanti J-26 XP centrifuge, and the supernatant was discarded. Cell pellets were stored at −20 °C.

Dh2$^{SC}$ and Dh10$^{SC}$ with an N-terminal $His_6$-SpyCatcher003 fusion and BMC-T1$^{ST}$ were purified by $Ni^{2+}$-NTA affinity chromatography followed by size-exclusion chromatography. Frozen cell pellets were thawed on ice and resuspended in 30 mL of lysis buffer (50 mM Tris pH 8, 300 mM NaCl, 10 mM imidazole, 5% v/v glycerol) with the addition of 1/2 Sigma protease inhibitor tablet and 200 µL of 2 mg/mL DNase I. Cells were chemically lysed by B-PER Bacterial Protein Extraction Reagent (Thermo Scientific) in 5:1 reagent to g cell pellet ratio with inverted mixing for 15 min. Cell lysate was clarified by centrifugation at 20,000 rpm for 1 h at 4 °C with a JA 20 rotor in a Beckman Coulter Avanti J-26 XP centrifuge. Supernatants were filtered using a 0.22 µm filter and transferred to clean tubes. Column chromatography was performed using an ÄKTA Start chromatography system (GE Healthcare), and clarified lysates were applied to a 5 mL HisTrap column (GE Healthcare) that was equilibrated with Buffer A (20 mM Tris–HCl pH 8, 500 mM NaCl). The column was washed with 6 column volumes (CV) of Buffer A and then subsequently washed with 6 CV of 98% Buffer A and 2% Buffer B (20 mM Tris–HCl pH 8, 500 mM NaCl, 500 mM imidazole). Protein was eluted over a 10 CV gradient from 2–100% Buffer B. Fractions containing the target protein were identified by SDS-PAGE analysis and were pooled and concentrated with a 10 kDa MWCO Amicon Ultra centrifugal filter (EMD Millipore). Using an ÄKTA Pure protein purification system, proteins were loaded onto a HiLoad 16/600 Superdex 200 pg (Sigma-Aldrich) size exclusion coulmn. Proteins were eluted in SEC buffer (50 mM Tris–HCl pH 8, 150 mM NaCl) using a flow rate of 1.0 mL/min. Purified proteins were concentrated, aliquoted and either flash-frozen and stored at -80 °C (BMC-T1$^{ST}$) or at 4 °C (Dh2$^{SC}$ and Dh10$^{SC}$).

BMC-H sheet purification was performed as described in (Range, Chiang et al., 2025). Frozen cell pellets from 1 L culture were resuspended in 30 mL of hexamer lysis buffer (50 mM Tris–HCl pH 8, 100 mM NaCl, 10 mM $MgCl_2$), 200 µL of 2 mg/mL DNase 1, and 100 µL of 10 mg/mL lysozyme. Cells were lysed by 3 passes through an Emulsiflex C3 Homogenizer (Avestin Inc) at 20,000 psi. 300 µL of Triton X-100 (1% v/v) was added to the lysate and incubated at room temperature with gentle agitation for 20 min on an orbital shaker. Insoluble material was separated by centrifugation for 20 min at 20,000 rpm with a JA 20 rotor in a Avanti J-26 XP centrifuge (Beckman Coulter). The supernatant was discarded, and the white pellet was resuspended in 30 mL of hexamer wash buffer (50 mM Tris–HCl pH 8, 100 mM NaCl, 10 mM $MgCl_2$, 1% v/v Triton X-100) without disturbing the brown cellular debris. The resuspension was transferred to a new centrifuge tube. The centrifugation/wash steps were repeated with 20–30 mL of hexamer wash buffer until the pellet was visibly more white and cellular debris was absent. The pellet was then washed once with 30 mL of hexamer lysis buffer to remove Triton X-100. The pellet was then resuspended in 10 mL of hexamer lysis buffer, aliquoted, pelleted, and stored at -80 °C as aliquots.

**Dh4 and Dh5:** Dh4$^{SC}$ and Dh5$^{SC}$ with an N-terminal $His_6$-SpyCatcher003 fusion and mixed BMC-T1-ST003 were heterologously expressed in *E. coli* BL21(DE3) and purified by $Ni^{2+}$-NTA affinity chromatography followed by SEC. Plasmids were transformed into *E. coli* BL21(DE3), and a 50 ml culture inoculated with single colony was grown overnight at 37 °C in LB containing 50 µg/mL kanamycin (100 µg/mL ampicillin for BMC-T1). One liter of LB supplemented with kanamycin (ampicillin for BMC-T1$^{ST}$) was inoculated with 1% (v/v) overnight culture and grown at 37 °C, 180 rpm until the $OD_{600}$ reached 0.6-0.8. Expression was induced with 100 ng/µl aTc (0.1 mM IPTG for BMC-T1), and cultures were grown overnight at 22 °C. Cells were harvested, flash-frozen, and stored at -80 °C. Cell pellets were resuspended in 40 mL TBS (50 mM Tris-HCl, pH 8.0, 150 mM NaCl) containing 20 mM imidazole and lysed using a homogenizer. After centrifugation at 20,000 × *g* for 45 min, the soluble fraction was applied to a 5 ml $Ni^{2+}$-NTA HiTrap column, washed with 20 mM imidazole and 300 mM NaCl, and eluted with 300 mM imidazole in TBS. Eluted proteins were concentrated and further purified by SEC on a Superdex™ 200 column equilibrated in TBS. Purified proteins were concentrated, aliquoted and either flash-frozen and stored at -80 °C 9(T1)  or at 4 °C  (Dh4 and Dh5).

BMC-H sheets were overexpressed by inoculating 1 L LB medium containing 50 µg/mL ampicilin with 1% (v/v) overnight culture and growing at 37 °C, 180 rpm to an $OD_{600}$ of 0.6-0.8. Expression was induced with 0.25 mM IPTG, and cultures were incubated overnight at 22 °C with shaking at 180 rpm. Cells were harvested, flash-frozen, and stored at -80 °C. BMC-H sheet purification was performed as described in our previous report (Range, Chiang et al., 2025). Frozen pellets were resuspended in 30 mL hexamer lysis buffer (50 mM Tris-HCl pH 8.0, 100 mM NaCl, 10 mM $MgCl_2$) supplemented with DNase I and lysozyme and lysed using a homogenizer. Triton X-100 was added to 1% (v/v), and lysates were incubated for 20 min at room temperature with gentle agitation. Insoluble white material enriched in BMC-H sheets formed on top of cell debris was collected after centrifugation at 20,000 rpm for 20 min, washed repeatedly with hexamer wash buffer (1% Triton X-100), followed by a final wash without detergent. The final pellet was resuspended in 10 mL hexamer lysis buffer, aliquoted, pelleted and stored at -80 °C as aliquots.

**Dh6 and Dh7:** Dh6$^{SC}$ and Dh7$^{SC}$ were expressed in *Escherichia coli* BL21(DE3). Transformed cells were initially grown overnight in LB medium supplemented with 50 µg $mL^{-1}$ kanamycin. The overnight culture was then diluted 1:100 (v/v) into fresh LB medium containing 50 µg $mL^{-1}$ kanamycin and incubated at 37 °C with shaking at 200 rpm. When the cell density reached an $OD_{600}$ of approximately 1.0, protein expression was induced by the addition of aTc to a final concentration of 100 ng $\mu L^{-1}$, followed by incubation at 20 °C with shaking at 200 rpm overnight.

Cells were harvested by centrifugation at 8,000 × *g* for 10 min at 4 °C, and the resulting pellets were resuspended in lysis buffer (20 mM Tris–HCl, 250 mM NaCl, 20 mM imidazole, pH 7.6). Cell lysis was performed using a French press. The lysate was clarified by centrifugation at 18,000 × *g* for 60 min at 4 °C, and the supernatant was collected and applied to a HisTALON gravity column (Takara) pre-equilibrated with lysis buffer. After sample loading, the resin was washed with lysis buffer to remove nonspecifically bound proteins. The target proteins were eluted using an elution buffer (20 mM Tris-HCl, 250 mM NaCl, 250 mM imidazole, pH 7.6). The elution fractions containing Dh6 or Dh7 were concentrated and exchanged into lysis buffer using

Amicon® Ultra centrifugal filters (30 kDa, MWCO) and then stored at 4 °C for further studies. Protein purity was analyzed by SDS–PAGE using 4–20% Mini-PROTEAN® TGX™ precast gels, and Precision Plus Protein™ Dual Color Standards (Bio-Rad).

The expression and purification of BMC-T1003 and BMC-H were performed as previously described. The purified BMC-T1003 was stored in lysis buffer at 4 °C.

**Dh11, Dh12 and Dh13:** Dh11$^{SC}$, Dh12$^{SC}$, and Dh13$^{SC}$ with a C-terminal SpyCatcher003-His$_6$ fusion and BMC-T1$^{ST-R}$ were heterologously expressed in *E. coli* BL21(DE3) and purified by $Ni^{2+}$-NTA affinity chromatography followed by size-exclusion chromatography. Plasmids were transformed into *E. coli* BL21(DE3), and single colonies were grown overnight at 37 °C in LB containing 50 µg/mL kanamycin (carbenicillin for BMC-T1). One liter of LB supplemented with kanamycin (carbenicillin for BMC-T1$^{ST-R}$) was inoculated with 1% (v/v) overnight culture and grown at 37 °C, 180 rpm until the $OD_{600}$ reached 0.6-0.8. Expression was induced with 100 ng/µL (0.3 mM IPTG for BMC-H), and cultures were grown overnight at 18 °C. Cells were harvested, flash-frozen, and stored at -80 °C. Cell pellets were resuspended in 40 mL TBS (50 mM Tris-HCl, pH 8.0, 150 mM NaCl) supplemented with protease inhibitors and lysed using a Constant Systems cell disruptor. After centrifugation at 20,000 × *g* for 20 min, the soluble fraction was applied to a $Ni^{2+}$-NTA column, washed with 20 mM imidazole, and eluted with 250 mM imidazole in TBS. Eluted proteins were concentrated and further purified by size-exclusion chromatography on a Superdex™ 200 column equilibrated in TBS containing 10% (v/v) glycerol. Purified proteins were concentrated, aliquoted, flash-frozen, and stored at -20 °C.

BMC-H sheets were overexpressed by inoculating 1 L LB medium containing 50 µg/mL carbenicillin with 1% (v/v) overnight culture and growing at 37 °C, 180 rpm to an $OD_{600}$ of 0.6-0.8. Expression was induced with 1 mM IPTG, and cultures were incubated overnight at 22 °C with shaking at 180 rpm. Cells were harvested, flash-frozen, and stored at -80 °C. BMC-H sheet purification was performed as described in (Range, Chiang et al., 2025). Frozen pellets were resuspended in 30 mL hexamer lysis buffer (50 mM Tris-HCl pH 8.0, 100 mM NaCl, 10 mM $MgCl_2$) supplemented with DNase I and lysozyme and lysed using a Constant Systems cell disruptor. Triton X-100 was added to 1% (v/v), and lysate was incubated for 20 min at room temperature with gentle agitation. After centrifugation at 20,000 rpm for 20 min, insoluble white material enriched in BMC-H sheets formed on top of cell debris was collected, washed repeatedly with hexamer wash buffer (1% Triton X-100), followed by a final wash without detergent. The purified pellet was resuspended in 10 mL hexamer lysis buffer and stored at -20 °C as aliquots.

**Dh14, Dh15 and Dh16:** For purification of BMC-T1$^{ST-R}$ and dehydrogenases, 2 L of LB were inoculated with 5 mL of preculture at $OD_{600}$ = 1.0 and grown to $OD_{600}$ = 0.8 before being induced with 100 mM IPTG or 100 ng/mL aTc (Millipore Sigma). Cells were harvested after 16-18 h of growth by centrifugation at 8000 x *g* for 10 min at 4 °C (Sorvall LYNX 6000, Thermo Fisher) and resuspended in 100 mL of 50 mM Tris pH 8.0, 50 mM NaCl, and 20 mM imidazole by vortexing. 200 µL of DNAse I (Millipore Sigma) and 0.5 tablet of SigmaFast protease inhibitor (Millipore

Sigma) were added. Resuspended cells were lysed by passage through a pre-cooled French press at 1,100 PSI twice. The lysate was clarified by centrifugation at 45,000 x *g* for 30 min at 4°C (Sorvall LYNX 6000, Thermo Fisher). The supernatant was removed and filtered through a 0.22 µm syringe filter. Proteins were purified using a ÄKTA pure fast protein liquid chromatography system (Cytiva) with a 5 mL HisTrap column (Cytiva) and eluted with 50 mM Tris pH 8.0, 50 mM NaCl and 300 mM imidazole. Proteins were then filtered using a 0.22 µm syringe filter before being further purified using a HiLoad Superdex 200pg SEC column (Cytiva) with buffer containing 50 mM Tris pH 8.0 and 200 mM NaCl to separate proteins from contaminants. Proteins were concentrated using 30 kDa Amicon Ultra 15 mL centrifugal filter units (Millipore) for use in *in vitro* assembly.

Purification of BMC-H was performed as previously described (Range, Chiang et al., 2025) with the following modifications. BMC-H sheets were resuspended in resuspension buffer containing 8 M urea before storage at –20 ˚C.

#### *3.2 In vitro* assembly

**Dh1:** Aliquoted BMC-H sheets were centrifuged to remove buffer, and the pellet was resuspended in 200 µL of 5 M urea. The suspension was incubated at room temperature with gentle rocking for 15 min, then 800 µL TBS (50 mM Tris-HCl, pH 8.0, 150 mM NaCl) was added followed by centrifugation at 17,000 × *g* for 5 min. The supernatant containing urea-solubilized BMC-H was collected, and protein concentration was estimated by measuring the absorbance at 280 nm using an appropriate buffer blank. Concentrations were calculated using the theoretical molar extinction coefficient of BMC-H at $A_{280}$. Concentrations of BMC-H, BMC-T1$^{ST}$ and Dh1$^{SC}$ were determined by measuring absorbance at 280 nm and by using their respective theoretical molar extinction coefficients (Dh1$^{SC}$, 53,860 $M^{-1}$ $cm^{-1}$; BMC-T1$^{ST}$, 15,595 $M^{-1}$ $cm^{-1}$; BMC-H, 2,980 $M^{-1}$ $cm^{-1}$).

*In vitro* assembly reactions (500 µL) were initiated by mixing BMC-T1$^{ST}$ and Dh1$^{SC}$ at a monomer ratio of 3:1 in TBS (50 mM Tris-HCl, 150 mM NaCl, pH 8.0) supplemented with 10% (v/v) glycerol, followed by incubation at room temperature for 60 min to allow SpyCatcher-SpyTag conjugation. BMC-H was then added at a BMC-H:BMC-T1$^{ST}$ tile ratio of 3:1. HT1 shell assembly occurred rapidly, and reaction mixtures were injected onto a Superdex™ 200 size-exclusion column for shell purification. Assembled shells eluted in the void volume, while unincorporated components eluted according to their molecular sizes. Void-volume fractions were collected. Assembly quality was assessed by dynamic light scattering (DLS).

**Dh2:** Aliquoted BMC-H sheet pellets were resuspended in 62.5 µL 8 M urea. The suspension was incubated at room temperature with gentle rocking for 1 h, then 937.5 µL of Tris-HCl pH 8 buffer (10 mM) was added. The final suspension was incubated at room temperature with gentle shaking overnight. Following incubation, suspension was centrifuged at 16,000 x *g* for 5 minutes. The supernatant containing urea solubilized BMC-H tiles was collected, and protein concentration was estimated by measuring absorbance at 280 nm using an appropriate buffer blank. Concentrations of BMC-H, BMC-T1$^{ST}$, and Dh2$^{SC}$ were determined by measuring absorbance at 280 nm and by using their theoretical molar extinction coefficient (Dh2$^{SC}$, 25,245 $M^{-1}$ $cm^{-1}$; BMC-T1$^{ST}$, 15,595 $M^{-1}$ $cm^{-1}$; BMC-H, 2,980 $M^{-1}$ $cm^{-1}$).

*In vitro* assembly reactions (1 mL) were initiated by mixing BMC-T1$^{ST}$ and Dh2$^{SC}$ at a monomer ratio of 1:1 in TBS (50 mM Tris-HCl, 150 mM NaCl, pH 8.0), followed by incubation at room temperature for 1 h to allow SpyTag-SpyCatcher conjugation. BMC-H was then added at a BMC-H:BMC-T1$^{ST}$ tile ratio of 3:1. HT1 shell assembly occurred rapidly, and reaction mixtures were injected onto a Superdex 200 Increase 10/300 GL by Cytiva size-exclusion column for shell purification. Assembled shells eluted in the void volume, while unincorporated components eluted according to their molecular sizes. Void-volume fractions were collected. Assembly quality was assessed by DLS.

**Dh4 and Dh5:** Aliquoted BMC-H sheets were centrifuged to remove buffer, and the pellet was resuspended in 200 µL 5 M urea. The suspension was incubated at room temperature with gentle rocking for 15 min, then 800 µL TBS (50 mM Tris-HCl, pH 8.0, 150 mM NaCl) was added followed by centrifugation at 17,000 × *g* for 5 min. The supernatant containing urea-solubilized BMC-H was collected, and protein concentration was estimated by measuring absorbance at 280 nm using an appropriate buffer blank. Concentrations were calculated using the theoretical molar extinction coefficient of BMC-H at $A_{280}$. Concentrations of BMC-H, BMC-T1$^{ST}$ and Dh4$^{SC}$ and Dh5$^{SC}$ were determined by measuring absorbance at 280 nm and by using their respective theoretical molar extinction coefficients (BMC-T1-SpyT, 15,595 $M^{-1}$ $cm^{-1}$; BMC-H, 2,980 $M^{-1}$ $cm^{-1}$).

*In vitro* assembly reactions (500 µL) were initiated by mixing BMC-T1$^{ST}$ and Dh4$^{SC}$ and Dh5$^{SC}$ at a monomer ratio of 3:1 in TBS (50 mM Tris-HCl, 150 mM NaCl, pH 8.0) supplemented with 10% (v/v) glycerol, followed by incubation at room temperature for 60 min to allow SpyCatcher-SpyTag conjugation. BMC-H was then added at a BMC-H:BMC-T1$^{ST}$ tile ratio of 3:1. HT1 shell assembly occurred rapidly, and reaction mixtures were injected onto a Superdex™ 200 size-exclusion column for shell purification. Assembled shells eluted in the void volume, while unincorporated components eluted according to their molecular sizes. Void-volume fractions were collected. Assembly quality was assessed by DLS.

**Dh6 and Dh7:** The conjugation of a T1 tile to Dh6$^{SC}$/Dh7$^{SC}$ was performed at a molar ratio of 1.5:1 and incubated on ice for 2 h. Then, H was added to the Dh-T1 mixture at a molar ratio of 3:1 (H: T) and incubated for 10 min. The mixed sample was loaded onto a Superdex 200 pg HiLoad 16/600 chromatography column (Cytiva) at a flow rate of 1 mL/min, using the lysis buffer as the elution buffer. The HT1-Dh6$^{SC}$/Dh7$^{SC}$ shell was eluted at around 40 min. Protein concentration was determined using a Nanodrop (Thermo Fisher, USA).

**Dh10:** Aliquoted BMC-H sheet pellets were resuspended in 100 µL 8 M urea. The suspension was incubated at room temperature with gentle rocking for 2 h, then 900 µL of Milli Q water was added. The final suspension was incubated at room temperature with gentle shaking overnight. Following incubation, suspension was centrifuged at 17,000 x *g* for 10 min. The supernatant containing urea solubilized BMC-H tiles was collected, and protein concentration was estimated by measuring absorbance at 280 nm using an appropriate buffer blank. Concentrations of BMC-H, BMC-T1$^{ST}$, and Dh10$^{SC}$ were determined by measuring absorbance at 280 nm and by using their theoretical molar extinction coefficient (Dh10$^{SC}$, 37,945 $M^{-1}$ $cm^{-1}$; BMC-T1$^{ST}$, 15,595 $M^{-1}$ $cm^{-1}$; BMC-H, 2,980 $M^{-1}$ $cm^{-1}$).

*In vitro* assembly reactions (1 mL) were initiated by mixing BMC-T1$^{ST}$ and Dh10$^{SC}$ at a monomer ratio of 1:1 in TBS (50 mM Tris-HCl, 150 mM NaCl, pH 8.0), followed by incubation at room temperature for 1 h to allow SpyTag-SpyCatcher conjugation. BMC-H was then added at a BMC-H:BMC-T1$^{ST}$ tile ratio of 3:1. HT1 shell assembly occurred rapidly, and reaction mixtures were injected onto a Superdex 200 Increase 10/300 GL by Cytiva size-exclusion column for shell purification. Assembled shells eluted in the void volume, while unincorporated components eluted according to their molecular sizes. Void-volume fractions were collected. Assembly quality was assessed by DLS.

**Dh11, Dh12, Dh13:** Aliquoted BMC-H sheets were centrifuged to remove buffer, and the pellet was resuspended in 500 µL TBS (pH 8.0) containing 1 M urea. The suspension was incubated at room temperature with gentle rocking for 15 min, followed by centrifugation at 17,000 × *g* for 5 min. The supernatant containing urea-solubilized BMC-H was collected, and protein concentration was estimated by measuring absorbance at 280 nm using an appropriate buffer blank. Concentrations were calculated using the theoretical molar extinction coefficient of BMC-H at $A_{280}$. Concentrations of Dh11$^{SC}$, Dh12$^{SC}$, Dh13$^{SC}$, and BMC-T1 were determined by measuring absorbance at 280 nm and by using their respective theoretical molar extinction coefficients (Dh11$^{SC}$, 38,975 $M^{-1}$ $cm^{-1}$; Dh12$^{SC}$, 45,840 $M^{-1}$ $cm^{-1}$; Dh13$^{SC}$, 43,890 $M^{-1}$ $cm^{-1}$; BMC-T1$^{ST\text{-}R}$, 15,595 $M^{-1}$ $cm^{-1}$; BMC-H, 2,920 $M^{-1}$ $cm^{-1}$).

*In vitro* assembly reactions (500 µL) were initiated by mixing BMC-T1 and the respective dehydrogenase at a monomer ratio of 3:2 in TBS (50 mM Tris-HCl, 250 mM NaCl, pH 8.0) supplemented with 10% (v/v) glycerol, followed by incubation at room temperature for 30 min to allow SpyCatcher-SpyTag conjugation. BMC-H was then added at a BMC-H:BMC-T1$^{ST\text{-}R}$ tile ratio of 3:1. HT1 shell assembly occurred rapidly, and reaction mixtures were clarified by high-speed centrifugation before injection onto a Superdex™ 200 size-exclusion column for shell purification. Assembled shells eluted in the void volume, while unincorporated components eluted according to their molecular sizes. Void-volume fractions were collected, and shell concentrations were determined using Pierce™ BCA Protein Assay Kits. Assembly quality was assessed by transmission electron microscopy (TEM) and DLS.

**Dh14:** *In vitro* assembly was performed as previously described (Range et al) with the following modifications: Glycerol was omitted from the reaction volume. For a 5 mL assembly 630 µL of 11.87 mg/mL BMC-H, 820 µL of 3.27 mg/mL BMC-T1$^{ST\text{-}R}$, and 1100 µL of 1.84 mg/mL Dh14$^{SC}$ was used. HT1-Dh14$^{SC}$ shells were separated from unassembled components using a HiLoad

Superdex 200 pg SEC column (Cytiva). Proteins were concentrated using 100 kDa Amicon Ultra 0.5 mL centrifugal filter units (Millipore) for use in TEM.

**Dh16:** Dh16$^{SC}$ was prepared as above but in 500 µL final volume: 63 µL of 11.87 mg/mL BMC-H, 82 µL of 3.27 mg/mL BMC-T1, and 17.4 µL of 8.48 mg/mL Dh16$^{SC}$. HT1-Dh16$^{SC}$ was separated using a Superdex 200 Increase 10/300 GL (Cytiva).

#### *3.3* Activity assays

**Dh1:** The catalytic activity of Dh1$^{SC}$, an aldehyde reductase from *E. coli*, was measured by continuously monitoring absorbance at 340 nm. Reactions were carried out at 22 °C and contained TBS (50 mM Tris-HCl, 150 mM NaCl, pH 8.0), 2 mM propionaldehyde, 0.4 mM NADPH and 10.7 µg of free Dh1$^{SC}$ or an equivalent amount of conjugated enzyme from an *in vitro* assembly reaction. Similarly, no-substrate control reactions were performed in the presence of HT1-Dh1$^{SC}$ and all other reaction components except the respective substrates. Specific activities, defined as activity per microgram of enzyme, for HT1-Dh1$^{SC}$ were calculated by assuming 100% encapsulation of Dh1$^{SC}$ within HT1 shells, based on the SEC profiles of shell assembly. Similarly, for Dh1$^{SC}$-BMC-T1 conjugates, activities were calculated per mg of only the enzyme present in the mixture, rather than the total protein amount.

**Dh2:** The catalytic activity of Dh2$^{SC}$, an alcohol dehydrogenase from *Zymononas mobilis,* was measured by continuously monitoring absorbance at 340 nm. Reactions were carried out at 22 °C and contained TBS buffer (50 mM Tris-HCl, 150 mM NaCl, pH 8.0), 20 mM ethanol, 0.5 mM $NAD^+$, and 2 µM of free Dh2$^{SC}$ or an equivalent amount of conjugated enzyme from an *in vitro* assembly reaction. Similarly, no-substrate control reactions were performed in the presence of HT1-Dh2$^{SC}$ and all other reaction components except the respective substrates. Specific activities, defined as activity per microgram of enzyme, for HT1-Dh2$^{SC}$ was calculated by normalizing absorbance at 340 nm of loaded shells to equivalent empty shells and calculating the difference in protein absorbance at 280 nm. This difference was attributed to Dh2$^{SC}$ encapsulated inside shells and used to calculate protein concentration. For Dh-BMC-T1 conjugates, activities were calculated per mg of only the enzyme present in the mixture, rather than the total protein amount.

**Dh4:** The catalytic activity of Dh4$^{SC}$, an aldehyde reductase from *Vibrio harveyi*, was measured by continuously monitoring absorbance at 340 nm. Reactions were carried out at 22 °C and contained TBS (50 mM Tris-HCl, 150 mM NaCl, pH 8.0), 2 mM butyraldehyde, 2 mM $NAD^+$, and 5 µg of free Dh4$^{SC}$ or an equivalent amount of conjugated enzyme from an *in vitro* assembly reaction. Similarly, no-substrate control reactions were performed in the presence of HT1-Dh4$^{SC}$ and all other reaction components except the respective substrates. Specific activities, defined as activity per mg of enzyme, for HT1-Dh4$^{SC}$ were calculated by assuming 100% encapsulation of Dh4$^{SC}$ within HT1 shells, based on the SEC profiles of shell assembly. Similarly, for Dh4-BMC-T1 conjugates, activities were calculated per mg of only the enzyme present in the mixture, rather than the total protein amount.

**Dh5:** The catalytic activity of Dh5$^{SC}$, a propionaldehyde dehydrogenase from *Clostridium phytofermentas*, was measured by continuously monitoring absorbance at 340 nm. Reactions were carried out at 22 °C and contained 100 mM phosphate buffer, pH 8.0, 4 mM propionaldehyde, 2 mM $NAD^+$, and 1 µg of free Dh5$^{SC}$ or an equivalent amount of conjugated enzyme from an *in vitro* assembly reaction. Similarly, no-substrate control reactions were performed in the presence of HT1-Dh5$^{SC}$ and all other reaction components except the respective substrates. Specific activities, defined as activity per mg of enzyme, for HT1-Dh5$^{SC}$ were calculated by assuming 100% encapsulation of Dh5$^{SC}$ within HT1 shells, based on the SEC profiles of shell assembly. Similarly, for Dh-BMC-T1 conjugates, activities were calculated per mg of only the enzyme present in the mixture, rather than the total protein amount.

**Dh6, Dh7:** Dh6$^{SC}$/Dh7$^{SC}$and HT1-Dh6$^{SC}$/HT1-Dh7$^{SC}$ kinetic activity assays using pyruvate were measured by monitoring the changes in NADH absorbance at 340 nm using an Agilent BioTek Epoch 2 Microplate Spectrophotometer in 96-well flat-bottom Costar microplates. Reactions were performed in a final volume of 200 µL per well using lysis buffer (20 mM Tris–HCl, 250 mM NaCl, 20 mM imidazole, pH 7.6) as the assay buffer. The reaction mixture contained NADH at a final concentration of 0.25 mM. Absorbance at 340 nm was recorded every 10 s at room temperature, and initial rates were determined from the linear portion of the $A_{340}$ time course.

For kinetic analysis, initial rates were plotted as a function of pyruvate concentration and fitted to the Michaelis–Menten equation to obtain $K_m$ and $V_{max}$ by nonlinear regression using Python. The specific activity per mg in HT1-Dh$^{SC}$ was calculated based on estimating the amount of Dh$^{SC}$ by the assembly ratio.

For activity comparison, enzymatic reactions were normalized based on the amount of dehydrogenase present in each sample. Free Dh6$^{SC}$ was assayed using 4 µg of purified enzyme. For Dh6$^{SC}$-T1 conjugates prepared at a 1:1.5 molar ratio, the total amount of conjugate added to the reaction was adjusted to ensure that the mixture contained 4 µg of Dh6$^{SC}$. Similarly, for HT1-Dh6$^{SC}$ assemblies, the amount of protein complex used in the assay was estimated to contain 4 µg of encapsulated Dh6$^{SC}$. An analogous normalization strategy was applied for Dh7-based samples, with 0.13 µg of Dh7$^{SC}$ used per reaction. All reactions were carried out in a total volume of 200 µL, and the reaction mixture and assay conditions were identical to those described in the kinetic activity assay.

**Dh10:** The catalytic activity of Dh10$^{SC}$, an L-leucine dehydrogenase from *Exiguobacterium sibiricum*, was measured by continuously monitoring absorbance at 340 nm. Reactions were carried out at 22 °C and contained 100 mM glycine buffer (pH 9.0), 100 mM leucine, 50 mM $NAD^+$, and 3.3 µg of free Dh10$^{SC}$ or an equivalent amount of conjugated enzyme from an *in vitro* assembly reaction. Similarly, no-substrate control reactions were performed in the presence of HT1-Dh10$^{SC}$ and all other reaction components except the respective substrates. Specific activities, defined as activity per microgram of enzyme, for HT1-Dh10$^{SC}$, was calculated by normalizing absorbance at 340 nm of loaded shells to equivalent empty shells and calculating the difference in protein absorbance at 280 nm. This difference was attributed to Dh10$^{SC}$ encapsulated inside shells and used to calculate protein concentration. For Dh10$^{SC}$-BMC-T1 conjugates, activities were calculated per mg of only the enzyme present in the mixture, rather than the total protein amount.

**Dh11, Dh12, Dh13:** The catalytic activity of Dh11$^{SC}$, an L-lactate dehydrogenase from *Clostridium thermocellum*, was measured by continuously monitoring absorbance at 340 nm. Reactions were carried out at 45 °C and contained 50 mM MOPS buffer (pH 7.0), 50 mM sodium pyruvate, 0.3 mM NADH, 1 mM fructose-1,6-bisphosphate, and either 10 µg of free Dh11$^{SC}$, 10 µg (enzyme equivalent) of Dh11$^{SC}$-BMC-T1$^{ST\text{-}R}$ conjugate, or 14 µg of HT1-Dh11 $^{SC}$. Activity assays for Dh12$^{SC}$ were performed using malic acid, oxaloacetate, pyruvate, or L-lactate as substrates with their corresponding cofactors. Assays were conducted over a pH range of 6.0 to 10.0 and temperatures ranging from 30 °C to 45 °C. No significant activity was observed under these conditions. The catalytic activity of Dh13$^{SC}$, a malate dehydrogenase from *Thermus thermophilus*, was measured similarly by continuously monitoring absorbance at 340 nm. Reactions were performed at 45 °C and contained 50 mM potassium phosphate buffer (pH 8.0), 4 mM malic acid, 5 mM NAD$^+$, and either 1 µg of free Dh13$^{SC}$, 1 µg (enzyme equivalent) of Dh13$^{SC}$-BMC-T1$^{ST\text{-}R}$ conjugate, or 5 µg of HT1-Dh13$^{SC}$. 'No-NAD$^+$/NADH control' reactions were performed in the presence of HT1-Dh$^{SC}$ and all other reaction components except the respective cofactors. Similarly, 'no-substrate control' reactions were performed in the presence of HT1-Dh$^{SC}$ and all other reaction components except the respective substrates. Specific activities, defined as activity per mg of enzyme, for HT1-Dh$^{SC}$ were calculated by assuming 100% encapsulation of Dh$^{SC}$ within HT1 shells, based on the SEC profiles of shell assembly. Similarly, for Dh$^{SC}$-BMC-T1$^{ST\text{-}R}$ conjugates, activities were calculated per mg of only the enzyme present in the mixture, rather than the total protein amount.

**Dh14:** The activity of Dh14$^{SC}$ was measured in a 96 well plate (Greiner Bio-One, 655101) with 46 ng of Dh14$^{SC}$, 10 mM NAD$^+$ (Sigma, 10127965001), 30 mM 2,3-butanediol (VWR, b06681-25g), and TBS 50/200 in a final volume of 100 µL. Absorbance at 340 nm was measured in a H1M plate reader (Biotek) for 1 h with measurements taken every min, shaking with maximal amplitude and minimal speed.

**Dh16:** GAPDH (Dh16$^{SC}$) activity was measured using the Glyceraldehyde-3-phosphate Dehydrogenase Activity Assay Kit (Colorimetric) (Abcam, ab204732) as described in the product manual. Protein samples were standardized for loading using the BCA assay described above, with 1,680 µg of GAPDH or 7,915 µg of HT1-Dh16$^{SC}$ added to each well.

#### *3.4* Thermal stability of free Dh$^{SC}$ and HT1-Dh$^{SC}$

Thermal stabilities of Dh6$^{SC}$, Dh7$^{SC}$, HT1-Dh6$^{SC}$, and HT1-Dh7$^{SC}$ were assessed by incubating protein samples at 30, 35, 40, 45, 50, and 55 °C in a metal heating block for 15 min. Following heat treatment, samples were immediately subjected to activity assays. The enzymatic activities measured at 30 °C were set to 100%, and residual activities at higher temperatures were expressed relative to this value. For thermal stability experiments with Dh14$^{SC}$, samples were prepared using 46 ng of Dh-14 diluted in TBS (50/200) pH 8.0 and incubated at the indicated temperatures (22, 37, 42, 47, 52, 57, 62, and 67 °C) for 1 h, followed by cooling at room temperature for 15 min. Samples were then transferred to a 96-well plate for activity assays. Thermal stability was assessed by measuring activity at the 10-min time point, with the activity at 22 °C set to 100%. Activities at higher temperatures (37, 42, 47, 52, 57, 62, and 67 °C) were expressed as a percentage of the activity at 22 °C.

#### *3.5* Lyophilization of Dh2$^{SC}$

250 µL of purified HT1-Dh2$^{SC}$ shells were frozen at -80 °C and dried at -50 °C on a Labconco FreeZone 2.5 L Freeze Dryer for 20 h. The dried proteins were resuspended in TBS buffer (50 mM Tris-HCl, 150 mM NaCl, pH 8.0) and spun at 10,000 x *g* for 2 min. Activity reactions were carried out at 22 °C and contained 2.2 µM of HT1-Dh2$^{SC}$ in TBS (50 mM Tris-HCl, 150 mM NaCl, pH 8.0), 20 mM ethanol, and 0.5 mM $NAD^+$. Specific activities, defined as activity per microgram of enzyme, were calculated by normalizing absorbance at 340 nm to equivalent empty shells and measuring the difference in protein absorbance at 280 nm. This difference was attributed to Dh2$^{SC}$ encapsulated inside shells and used to calculate Dh2$^{SC}$ concentration inside the shell. Specific activities of lyophilized samples and equivalent untreated samples were used to calculate relative enzyme activity.

#### *3.6* Storage stability of Dh$^{SC}$ and HT1-Dh$^{SC}$ at 4 °C and at RT

The stability and longevity of the free and shell-encapsulated enzymes, Dh4$^{SC}$, HT1-Dh4$^{SC}$, Dh5$^{SC}$, and HT1-Dh5$^{SC}$, were compared by monitoring residual enzymatic activity over 6 weeks. Samples were stored at 4 °C and room temperature (approximately 22–25 °C). To prevent microbial contamination, 0.02% (w/v) sodium azide was added to all preparations. Residual activities were assessed every 7 days by measuring the absorbance decrease of NADH at 340 nm (corresponding to $NAD^+$ production in the enzymatic reaction; $\varepsilon = 6{,}220\ M^{-1}\ cm^{-1}$). Aliquots were withdrawn from each storage condition, and activity was determined using the standard spectrophotometric assay for the respective dehydrogenase enzymes.

#### *3.7* Statistical analysis

All results are presented as mean ± SD from at least three independent replicates (unless specifically stated). Statistical analyses were performed using GraphPad Prism version 9.0 (San Diego, CA, USA). Differences among groups were assessed using two-way analysis of variance (ANOVA). A p value of $< 0.05$ was considered statistically significant.

### 4. Supplementary figures

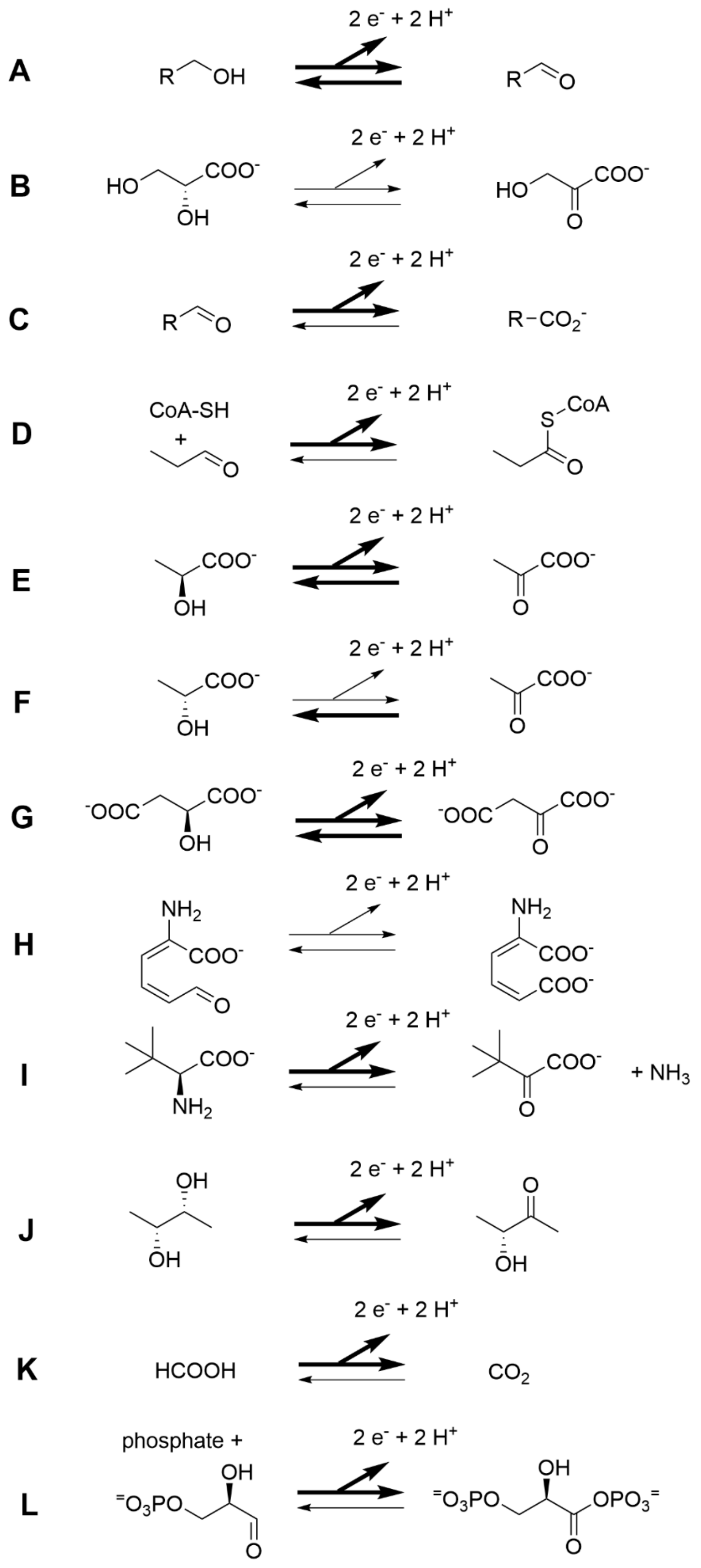

**Fig. S1. Oxidoreduction reactions catalyzed by the selected dehydrogenases.** Reactions catalyzed by the often-reversible enzymes are shown with the substrate being oxidized. Bold arrows indicate the direction of the reactions that were assayed to measure the activities of the corresponding enzymes. Two bold arrows indicate that one enzyme catalyzing the reaction was assayed in one direction (e.g., Dh1 in A), while another enzyme of the same class (e.g., Dh2 in A) was assayed in the opposite direction. **A)** Dh1: aldehyde reductase, Dh2: alcohol dehydrogenase, **B)** Dh3: glycerate dehydrogenase, **C)** Dh4: acetaldehyde dehydrogenase, **D)** Dh5: propionaldehyde dehydrogenase, **E)** Dh6 and Dh11: L-lactate dehydrogenase, **F)** Dh7: D-lactate dehydrogenase, **G)** Dh8, Dh12 and Dh13: malate dehydrogenase, **H)** Dh9: aminomuconate-semialdehyde dehydrogenase, **I)** Dh10: leucine dehydrogenase, **J)** Dh14: butanediol dehydrogenase, **K)** Dh15: formate dehydrogenase, **L)** Dh16: glyceraldehyde 3-phosphate dehydrogenase.

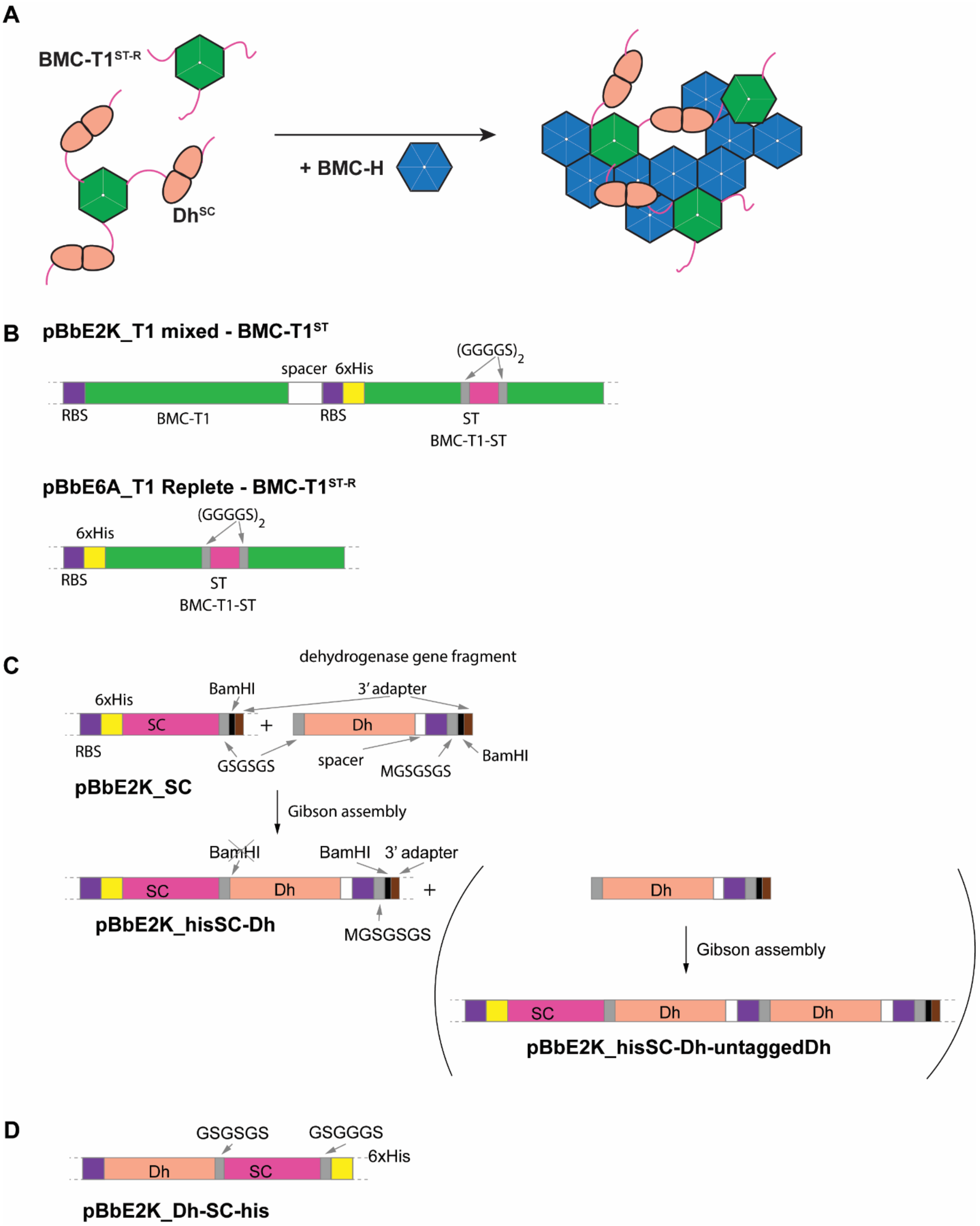


**Fig. S2. Construct considerations and cloning strategy. A)** Multimeric cargo with SpyCatcher (SC) can potentially interact with multiple BMC-T1$^{ST}$ (where ST is used for SpyTag) and hinder shell assembly when H subunits are added after conjugation. When all three monomers in the trimer have SpyTag fusions, they are designated as "replete" and denoted BMC-T1$^{ST-R}$. **B)** T1 construct for expressing mixed trimers containing both ST- and His$_6$-tagged copies as well as native copies of T1 (denoted as BMC-T1$^{ST}$). T1 construct for expressing only ST- and His$_6$-tagged copy (Tefft et al., 2026) (denoted as BMC-T1$^{ST-R}$) **C)** Cloning scheme for N-terminally His$_6$-SC-tagged Dh. In the first step the Dh gene fragment is inserted by Gibson cloning into the pBbE2K_SC vector cut with BamHI. The Gibson step eliminates the BamHI site, but the Dh fragment contains another BamHI site that can again accept the same Dh gene fragment to obtain a vector expressing His$_6$-SC-tagged Dh as well as untagged Dh. Steps shown in brackets were not performed in this work but could be useful for encapsulating proteins that form higher-order oligomers. **D)** C-terminally tagged SC-Dh gene fragments were cloned directly into pBbE2K.

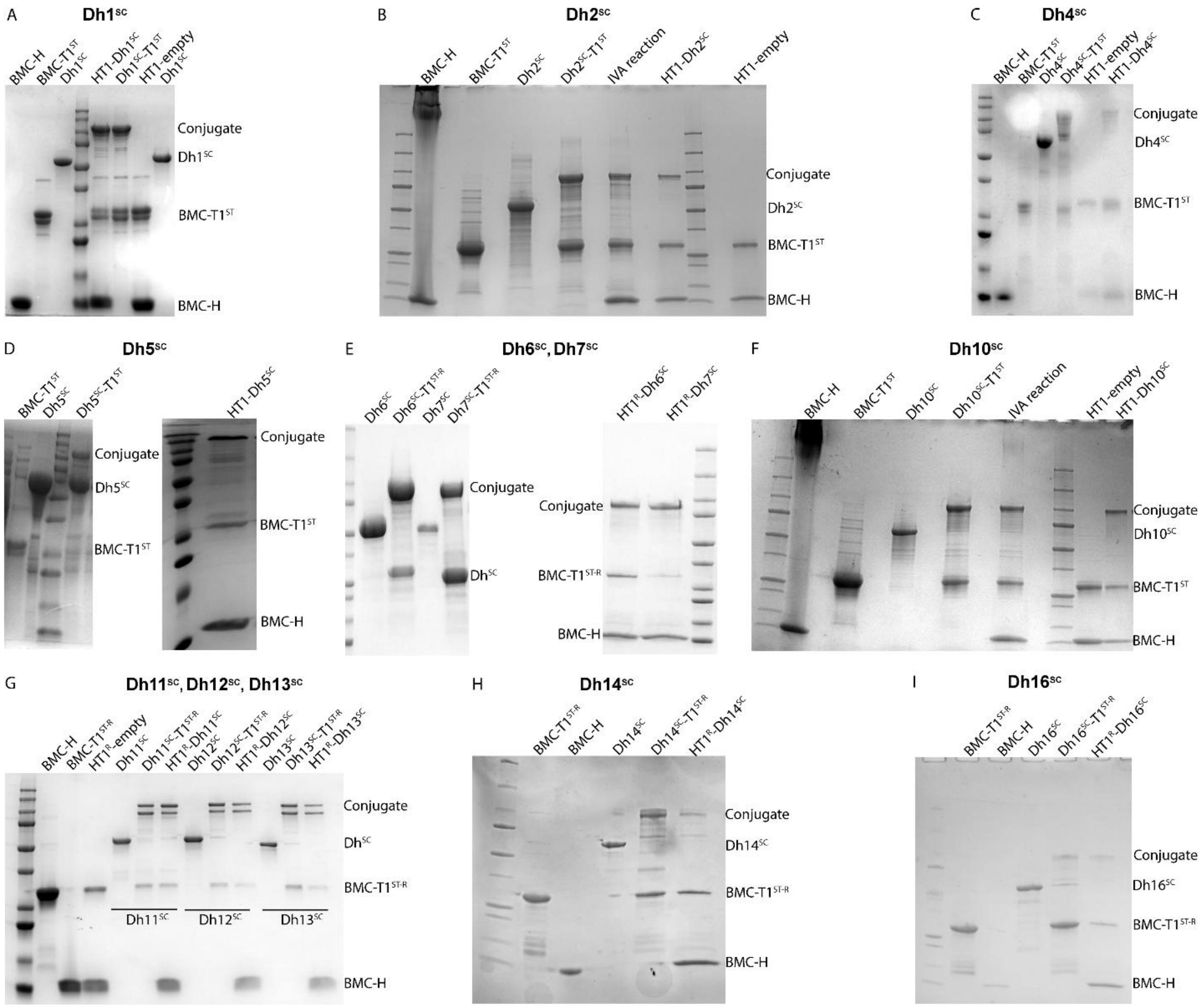


**Fig. S3. SDS-PAGE gels of HT1 shell components, cargo enzymes and their assembly.** Purification and HT1 shell assembly were performed in multiple laboratories across two institutions. SDS-PAGE gels show the purification of shell components (BMC-H and BMC-

T1$^{ST}$/BMC-T1$^{ST-R}$), Dh$^{SC}$, conjugation between Dh$^{SC}$ and BMC-T1$^{ST}$ (or BMC-T1$^{ST-R}$), and the assembly of both Dh$^{SC}$ loaded and empty HT1 shells by different researchers. Dh$^{SC}$ corresponds to Dh fused to SC; BMC-T1$^{ST}$ corresponds to BMC-T1 tiles with ST in at least one of the monomers and BMC-T1$^{ST-R}$ corresponds to BMC-T1 tiles with ST in all the monomers. Lanes in each gel are labeled as follows: **BMC-H**, purified and urea-reconstituted BMC-H sheets; **BMC-T1$^{ST}$/BMC-T1$^{ST-R}$**, purified BMC-T1 tiles; **Dh$^{SC}$**, purified dehydrogenases (with enzyme numbers corresponding to the nomenclature in Table 1); **Dh$^{SC}$-T1$^{ST}$/Dh$^{SC}$-T1$^{ST-R}$**, conjugation reaction showing covalent linkage between BMC-T1$^{ST}$/BMC-T1$^{ST-R}$ and Dh$^{SC}$, indicated by the appearance of a higher molecular-weight band (~100 kDa) corresponding to the combined mass of BMC-T1$^{ST}$/BMC-T1$^{ST-R}$ and Dh$^{SC}$; **IVA reaction**, *in vitro* assembly reaction containing Dh$^{SC}$, BMC-H, and BMC-T1 tiles prior to purification by SEC; **HT1-empty**, unloaded HT1 shells lacking encapsulated cargo; and **HT1-Dh$^{SC}$/HT1$^{R}$-Dh$^{SC}$**, HT1/HT1$^{R}$ shells encapsulating Dh$^{SC}$. Major protein bands are labeled on the right side of the gel according to their expected molecular weights, including free Dh$^{SC}$ (unconjugated dehydrogenase), the trimer-dehydrogenase conjugate, unconjugated BMC-T1$^{ST}$ /BMC-T1$^{ST-R}$, and BMC-H.

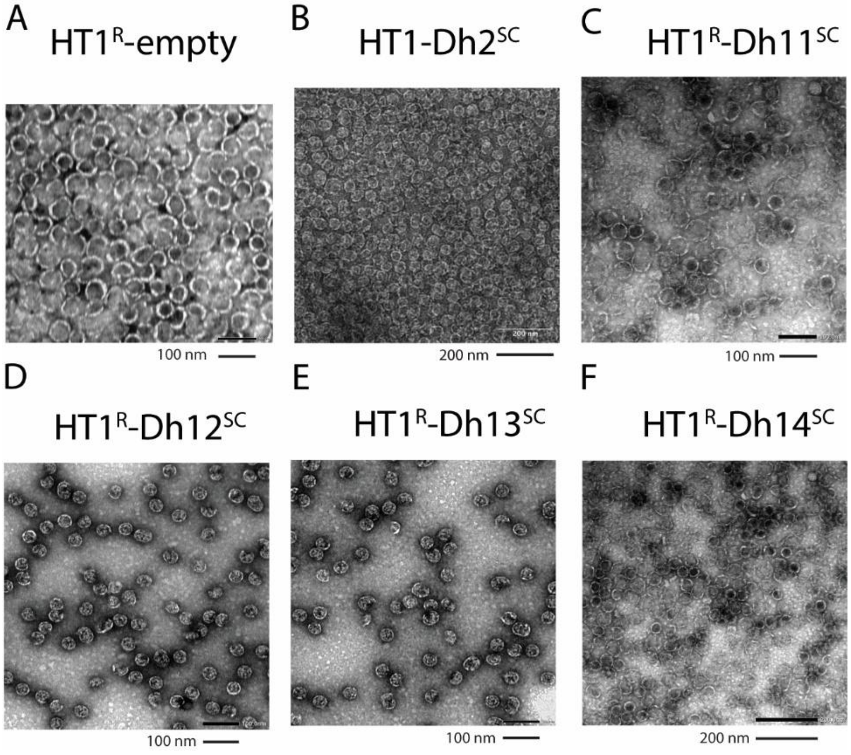


**Fig. S4**: **Transmission electron microscopy (TEM) images of empty and enzyme-loaded HT1 shells**. HT1 shells were assembled *in vitro*, purified by **SEC** to remove unassembled components, and subsequently imaged by TEM. The images show the formation of uniformly sized shell structures. HT1$^{R}$ denotes shells assembled using BMC-T1$^{ST-R}$. Scale bars differ between panels, as images were acquired by multiple researchers in different laboratories.

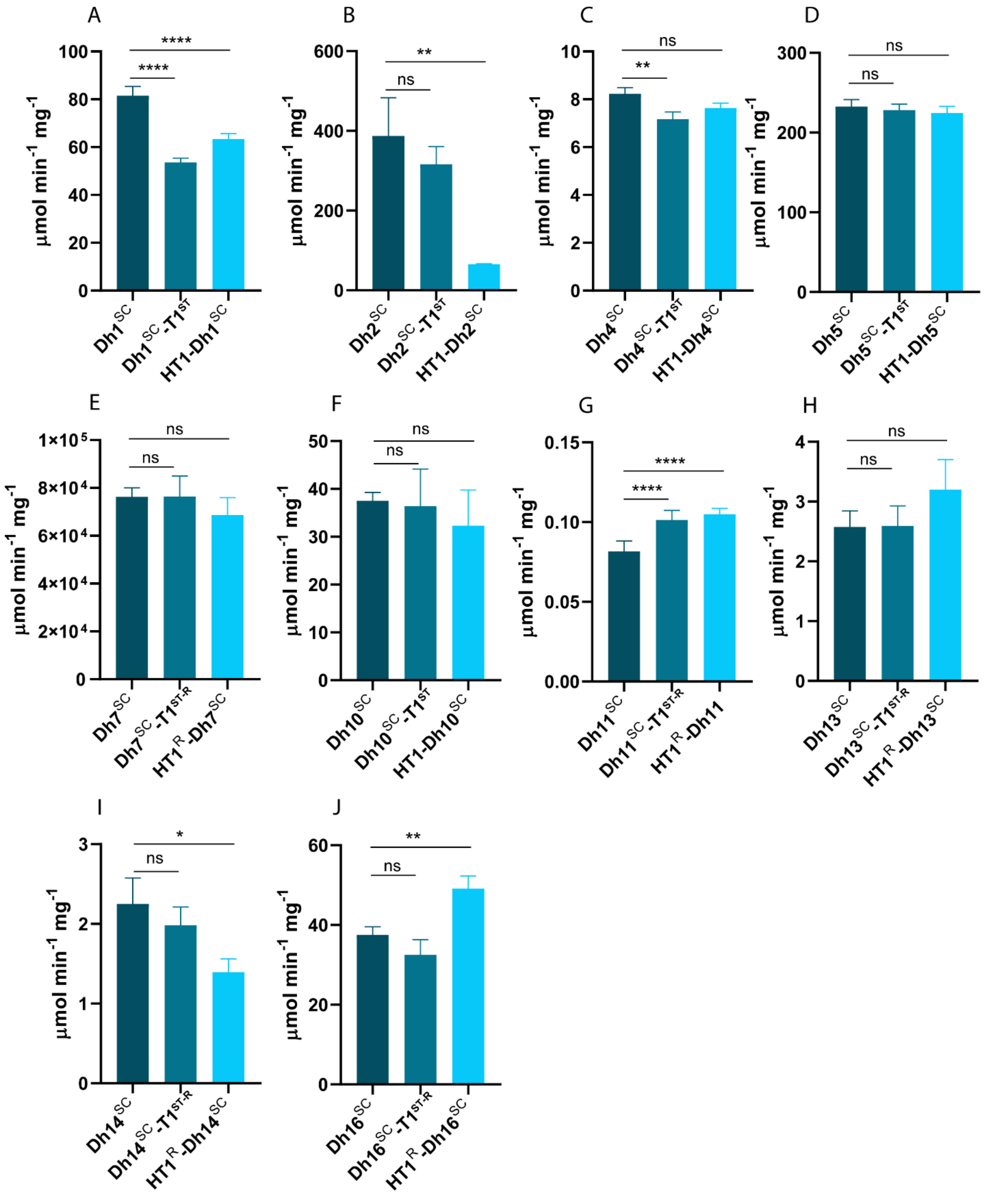


**Fig. S5: Enzyme specific differences in the catalytic activity between free SC-tagged dehydrogenases, conjugated dehydrogenases, and encapsulated dehydrogenases.** Enzyme activity per mg for conjugated enzymes was calculated based on the enzyme concentration in the conjugation reaction mixture. For encapsulated enzymes, it was calculated assuming complete encapsulation, as inferred from SEC profiles. Conjugation and encapsulation affect the catalytic activities of the dehydrogenases differently. $Dh^{SC}$, SC tagged dehydrogenases; $Dh^{SC}$-$T1^{ST}$ ($Dh^{SC}$-$T1^{ST\text{-}R}$), $Dh^{SC}$ conjugated to BMC-$T1^{ST}$(BMC-$T1^{ST}$); $HT1^{R}$, denotes shells assembled using BMC-$T1^{ST\text{-}R}$. Bars represent the mean, and error bars represent the standard deviation from at least three independent measurements. Statistical analysis was performed using one-way ANOVA in GraphPad Prism (ns $P > 0.05$; $^{*}P < 0.05$; $^{**}P < 0.005$; $^{****}P < 0.0001$).

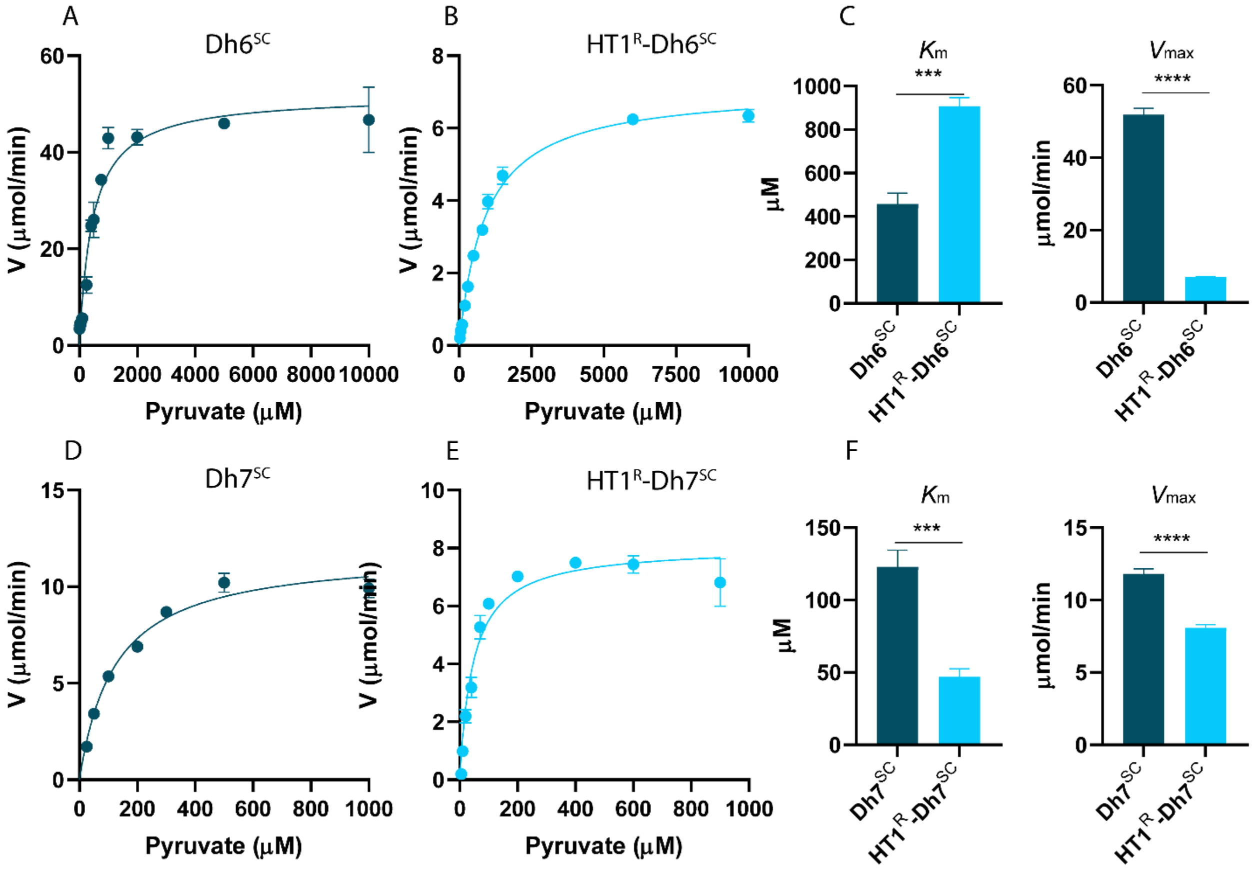


**Fig. S6: Effects of encapsulation of $Dh6^{SC}$ and $Dh7^{SC}$ in HT1 shells on enzyme kinetics. A)** Michaelis-Menten fit of free $Dh6^{SC}$, **B)** Michaelis-Menten fit of encapsulated $HT1^{R}$-$Dh6^{SC}$, **C)** Comparison of kinetic parameters ($K_m$ and $V_{max}$) for free and encapsulated $Dh6^{SC}$, highlighting a substantial increase in $K_m$ and decrease in $V_{max}$ upon encapsulation, indicative of stronger kinetic penalties, **D)** Michaelis-Menten fit of free $Dh7^{SC}$, **E)** Michaelis-Menten fit of encapsulated $HT1^{R}$-$Dh7^{SC}$, **F)** Comparison of kinetic parameters ($K_m$ and $V_{max}$) for free and encapsulated $Dh7^{SC}$, showing a pronounced decrease in $K_m$ with only a moderate reduction in $V_{max}$, reflecting enhanced apparent substrate affinity and favorable enzyme-shell compatibility. Bars represent the mean, and error bars represent the standard deviation from three independent measurements. Statistical analysis was performed using an unpaired t-test in GraphPad Prism ($^{***}P < 0.0005$; $^{****}P < 0.0001$).

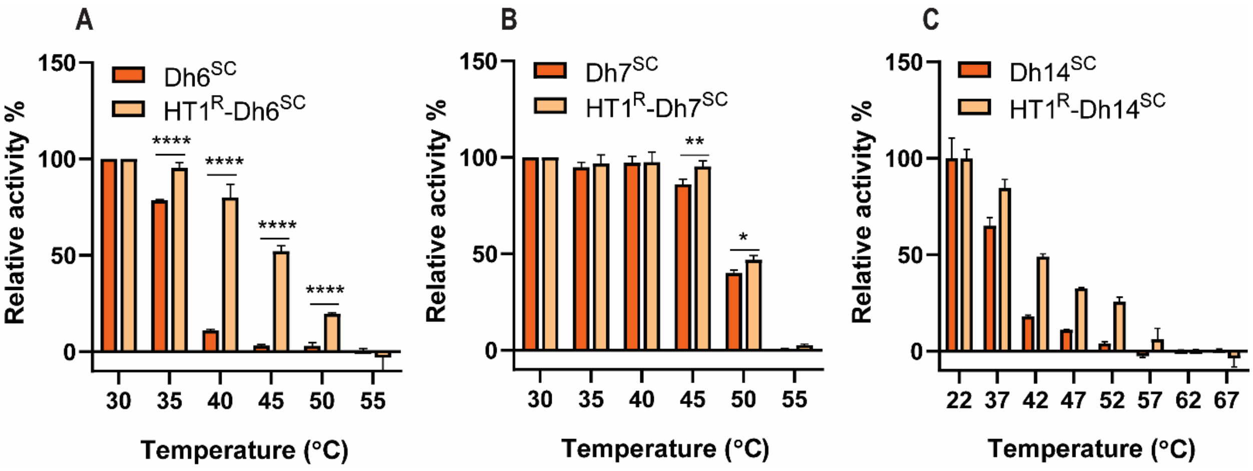


**Fig. S7**: **Effect of encapsulation of dehydrogenases in HT1$^R$ shells on their thermal stability.** **A)** Thermal stability of Dh6$^{SC}$ before and after encapsulation in HT1$^R$ shells. HT1$^R$-Dh6$^{SC}$ retained significantly greater residual activity at elevated temperatures (40-50 °C) compared to the free SC-tagged enzyme, indicating that shell encapsulation confers protection against thermal inactivation. **B)** Thermal stability of Dh7$^{SC}$ before and after encapsulation. Although Dh7$^{SC}$ is intrinsically more thermally stable than Dh6$^{SC}$, HT1$^R$-Dh7$^{SC}$ retained slightly greater relative activity than free Dh7$^{SC}$ at elevated temperatures, but both completely lost activity at 50 °C. **C)** Thermal stability of Dh14$^{SC}$ before and after encapsulation in HT1$^R$ shells. Stability conferred by encapsulation is intermediate for Dh14$^{SC}$. Bars represent the mean, and error bars represent the standard deviation from three independent measurements for A and B. Statistical analysis was performed using an unpaired t-test in GraphPad Prism (***$P < 0.0005$; ****$P < 0.0001$). Error bars in panel C represent the range from two independent measurements.

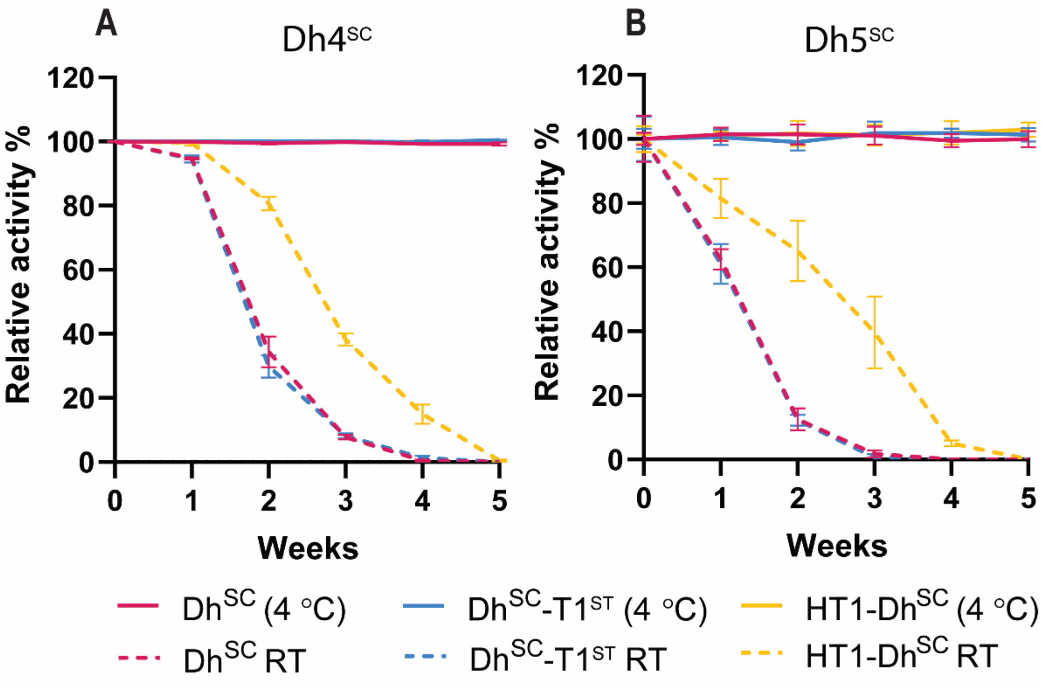


**Fig. S8: Effect of storage conditions on the catalytic activities of free SC-tagged, conjugated, and encapsulated Dh$^{SC}$. A)** Dh4$^{SC}$, Dh4$^{SC}$-T1$^{ST}$, and encapsulated HT1-Dh4$^{SC}$ stored at 4 °C (solid lines) did not show significant changes in activity over a period of 5 weeks, whereas encapsulated HT1-Dh4$^{SC}$ consistently exhibited greater activity when stored at room temperature (dotted line, RT). **B)** Similarly, Dh5$^{SC}$, Dh5$^{SC}$-T1, and encapsulated HT1-Dh5$^{SC}$ did not show significant changes in activity when stored at 4 °C (solid lines), while encapsulated HT1-Dh5$^{SC}$ retained greater activity when stored at RT (dashed line).

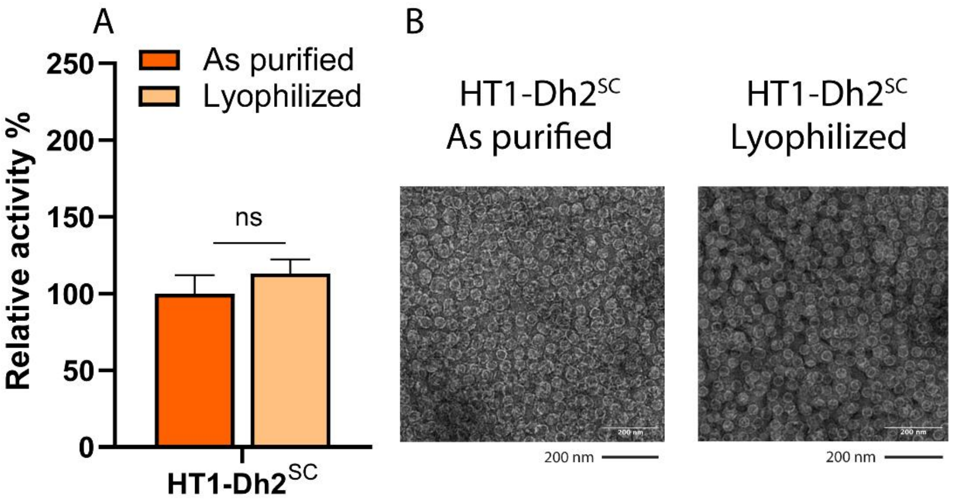


**Fig. S9: Effect of lyophilization on the catalytic activity and structural integrity of encapsulated HT1-Dh2$^{SC}$. A)** HT1-Dh2$^{SC}$ tolerated lyophilization well with the activity retained after reconstitution in buffer, **B)** TEM image showing that lyophilization does not affect the integrity of HT1-Dh2$^{SC}$ shells. Encapsulated HT1-Dh2$^{SC}$ (as purified) refers to SEC-purified shells stored at 4 °C, whereas HT1-Dh2$^{SC}$ (lyophilized) refers to SEC-purified shells that were lyophilized and subsequently reconstituted in TBS (50 mM Tris-HCl, 150 mM NaCl, pH 8.0). Bars in panel A represent the mean and error bars represent the standard deviation from three independent measurements. Statistical analysis was performed using two-way ANNOVA in GraphPad Prism (ns, P > 0.05).

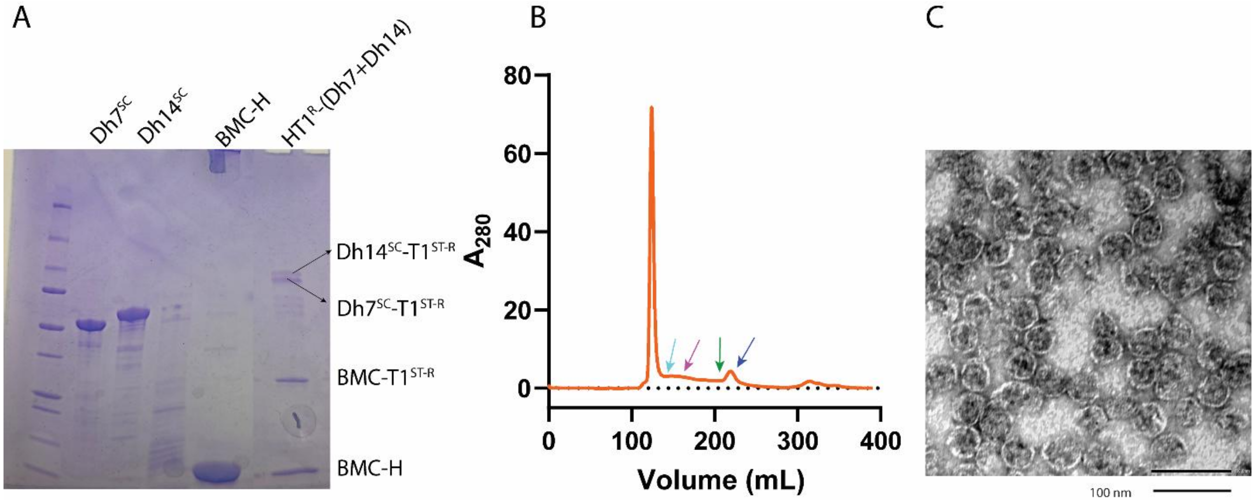


**Fig. S10: Encapsulation of both Dh7$^{SC}$ and Dh14$^{SC}$ into HT1$^{R}$ shells. A)** SDS-PAGE showing purified Dh7$^{SC}$, Dh14$^{SC}$, BMC-H, T1-conjugated enzymes and HT1$^{R}$-(Dh7+Dh14), **B)** Size exclusion chromatography showing assembly of Dh7$^{SC}$ and Dh14$^{SC}$ into HT1$^{R}$ shells. The presence of a single dominant peak in the void volume, with no significant peaks corresponding to free Dh$^{SC}$ (marked as ↓), BMC-T1$^{ST}$ or BMC-T1$^{ST-R}$ (marked as ↓), BMC-H (marked as ↓), or Dh-trimer conjugates (marked as ↓), indicates efficient assembly and near-complete incorporation of cargo enzymes. **C)** TEM of HT1$^{R}$-(Dh7$^{SC}$+Dh14$^{SC}$) shells showing well-formed intact shells.
